# Adipose GPD2 Couples Glycolytic Signals to Epigenetic Control of Metabolic Homeostasis

**DOI:** 10.64898/2026.07.19.739432

**Authors:** Yi-jie Shi, Meng Ding, Lin-xi Xiao, Ge Zhang, Xin Dou, Hong-yu Xu, Yi-fan Xia, Yi-fan Ke, Ze-yi Man, Jin-yi Xia, Lan-yue Zhang, Wei-jie Shan, Shu-wen Qian, Yan Tang, Qi-qun Tang, Yang Liu

## Abstract

The glycerol-3-phosphate shuttle (G3PS) is a classic reducing equivalent transferring system, yet its role in adipose metabolism remains poorly understood. Here, we identify mitochondrial glycerol-3-phosphate dehydrogenase (GPD2), the core enzyme of G3PS, as a critical orchestrator of adipose thermogenesis and systemic metabolic homeostasis. Adipocyte-specific GPD2 knockout mice exhibited impaired thermogenic capacity and energy expenditure, rendering them susceptible to obesity and metabolic dysfunction. Mechanistically, GPD2 deficiency elevated cytosolic NADH, which suppressed glycolysis and decreased the levels of key metabolites, such as acetyl-CoA, thereby diminishing H3K27ac at thermogenic gene loci and silencing these genes. NADH depletion with an NADH oxidase rescued both glycolysis and thermogenic gene expression. In support of this mechanism, we defined a previously undescribed enhancer of *Ucp1* whose H3K27ac was tightly controlled by GPD2. Remarkably, replenishing the acetyl-CoA pool or directly restoring H3K27ac at the *Ucp1* enhancer reversed UCP1 expression in GPD2-deficient adipocytes. Importantly, we find that human adipose GPD2 expression is inversely correlated with BMI, WHR, and HOMA-IR, underscoring its clinical relevance. Collectively, our findings establish GPD2 as an essential node in the metabolic-epigenetic axis that governs thermogenic activation and energy balance.

## Introduction

The global rise in obesity and its associated metabolic diseases highlights the need to elucidate the underlying pathological mechanisms. Adipose tissue dysfunction is a major driver of this pathophysiology^1–3^. Unlike energy-storing white adipocytes, thermogenic adipocytes, which comprise brown and beige fat, possess the remarkable ability to dissipate chemical energy as heat via uncoupling protein 1 (UCP1)-dependent or -independent non-shivering thermogenesis^4, 5^. This capacity positions the activation of thermogenic fat as a promising strategy to combat metabolic diseases^6^. Indeed, studies in human have shown that the absence of brown adipose tissue (BAT) is independently correlated with a high incidence of metabolic disorders^7^. Therefore, a deeper understanding of the mechanisms governing thermogenic adipocyte function holds great physiological and clinical significance.

Considerable progress has been made in deciphering the transcriptional control of thermogenic activation. Core regulators such as peroxisome proliferator activated receptor gamma (PPARγ), PR domain containing 16 (PRDM16), and PPARγ coactivator 1α (PGC1α), activated by upstream pathways including β-adrenergic signaling, are considered the principal orchestrators of thermogenic gene expression^8, 9^. While this perspective has successfully identified the terminal effectors of thermogenesis, it offers limited insight into the upstream metabolic signals that govern this energy-dissipating process. A growing body of evidence now indicates that cellular metabolic state is not merely an outcome of transcriptional regulation, but an active determinant of gene expression^10^. Key metabolites including lactate, α-ketoglutarate (α-KG) and acetyl-CoA serve as essential substrates or cofactors for epigenetic modifying enzymes, thereby functioning as critical links that couple metabolic state to stable changes in chromatin architecture and transcriptional landscape^11, 12^.

Mitochondrial glycerol 3-phosphate dehydrogenase (mGPDH, also known as GPD2) is a core component of the glycerol-3-phosphate shuttle (G3PS)^13^. Together with the malate-aspartate shuttle (MAS), the G3PS constitutes one of the primary systems for transferring reducing equivalents from cytosolic NADH into the mitochondria, thereby linking cytoplasmic glycolysis with mitochondrial oxidative phosphorylation^13^. As the rate-limiting enzyme within G3PS, GPD2 catalyzes the irreversible oxidation of glycerol-3-phosphate (G3P) to dihydroxyacetone phosphate (DHAP), concurrently transferring electrons via its FAD cofactor directly into the mitochondrial electron transport chain^13^. Moreover, the interconversion between G3P and DHAP regulates NAD^+^/NADH redox balance and serves as a critical metabolic hub linking glycolytic and lipid biosynthesis^13, 14^. Accumulating evidence has established GPD2 as a key regulator in diverse physiopathological processes, including cancer progression^13^, hepatic steatosis^15^, skeletal muscle regeneration^16^, diabetic kidney disease^17^, macrophage inflammatory responses^18^, and ferroptosis^19^. Thus, GPD2 functions as both a metabolic integrator and a regulator of mitochondrial bioenergetics and redox balance.

Our previous unbiased screen for regulatory factors in thermogenic adipocytes revealed a spectrum of key modulators, including CDO1^20^, CHCHD10^21^, and CLCF1^22^. Among these candidates, *Gpd2* was one of the most significantly upregulated genes during thermogenic activation. Here, we identified GPD2 as a metabolic checkpoint that controlled thermogenic capacity through epigenetic reprogramming. GPD2 deficiency inhibited glycolysis and reduced acetyl-CoA levels, thereby decreasing the enrichment of histone H3 lysine 27 acetylation (H3K27ac) at thermogenic loci and repressing their expression. Consequently, adipocyte-specific *Gpd2* knockout (GPD2^AKO^) mice exhibited impaired thermogenesis, increased weight gain, and metabolic disorders.

## Results

### Identification of adipose GPD2 as a signature of metabolic homeostasis

We first examined GPD2 expression across distinct adipose depots. GPD2 was detectable in inguinal white adipose tissue (iWAT), epididymal white adipose tissue (eWAT), and BAT, with the highest levels observed in BAT (Fig. 1A). In all three adipose tissues, *Gpd2* expression was significantly higher in mature adipocytes than in the stromal vascular fraction (SVF, Fig. 1B). Consistently, GPD2 levels increased progressively during brown and white adipocyte differentiation (Fig. 1C and D). At both the transcript and protein levels, GPD2 expression was markedly upregulated in response to cold exposure compared with thermoneutral (TN) conditions (Fig. 1E and F). Furthermore, treatment with β3-adrenergic receptor agonist CL316,243, which mimics cold stimulation to activate thermogenesis and energy metabolism, similarly promoted GPD2 expression across all three adipose depots (Fig. 1G).

**Fig. 1.**
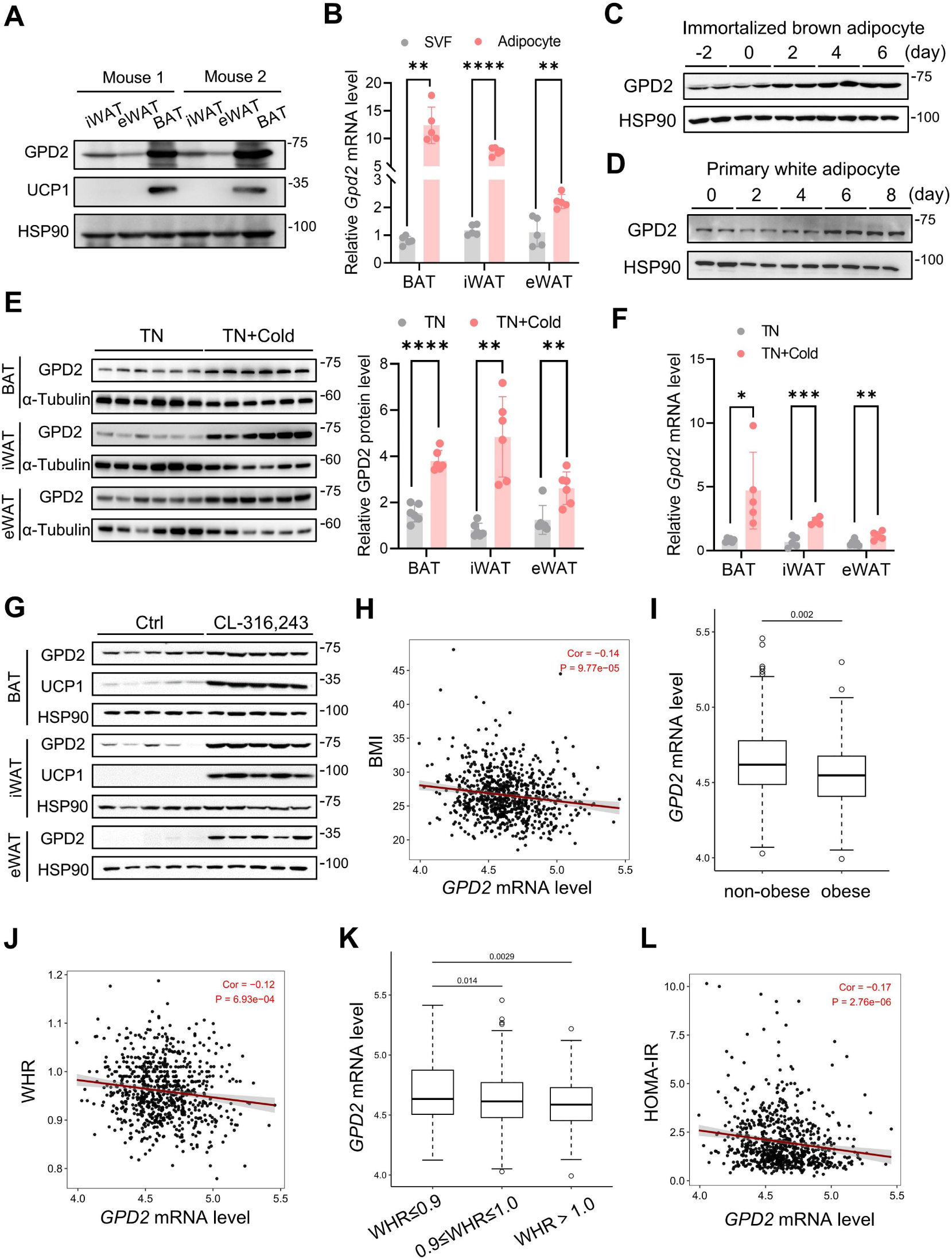
Adipose GPD2 is positively associated with thermogenic capacity. (A) GPD2 protein levels in BAT, iWAT and eWAT from 8-week-old male mice (n = 2). (B) *Gpd2* mRNA levels in mature adipocytes and SVF isolated from BAT, iWAT and eWAT of 8-week-old male mice (n = 5). Data were analyzed using two-tailed unpaired Student’s t-test (eWAT) or with Welch’s correction (BAT and iWAT). (C-D) GPD2 protein levels during adipogenic differentiation of immortalized brown preadipocytes (C) and primary WAT-derived stem cells (D). (E-F) Eight-week-old male mice were housed at thermoneutrality (TN, 30°C) for 2 weeks as controls or transferred from TN to 4°C for 72 h for cold exposure. Western blot showing GPD2 protein levels in three adipose tissues. Data were analyzed using two-tailed unpaired Student’s t-test (eWAT and BAT) or with Welch’s correction (iWAT). (F) qPCR data showing *Gpd2* mRNA levels in three adipose tissues (n = 4-6). Data were analyzed using two-tailed unpaired Student’s t-test (iWAT and eWAT) or with Welch’s correction (BAT). (G) GPD2 protein levels in BAT, iWAT, and eWAT from 8-week-old male mice administered with CL316,243 (1 mg/kg/day) for 7 consecutive days (n = 5). (H-L) Data were obtained from the GEO database (GSE70353). In human subcutaneous adipose tissue samples (n = 770), correlations between *GPD2* mRNA expression and BMI (H), WHR (J), and HOMA-IR (L) were analyzed. Significance was assessed using Spearman’s rank correlation test. Box plots show *GPD2* mRNA expression in abdominal subcutaneous adipose tissue between obese and non-obese individuals (I) and across WHR groups (K). P-values were calculated using the Mann-Whitney test. *, p < 0.05; **, p < 0.01; ***, p < 0.001.

Next, we assessed GPD2 expression under obese conditions, in which adipocyte energy metabolism is known to be impaired. In human adipose tissue, GPD2 expression was inversely correlated with body mass index (BMI), showing a pronounced reduction in individuals with obesity (Fig. 1H and I). Consistently, GPD2 expression also exhibited a significant negative correlation with waist-to-hip ratio (WHR) and the homeostasis model assessment of insulin resistance (HOMA-IR, Fig. 1J-L). Collectively, these data demonstrated that adipose GPD2 was positively correlated with metabolic homeostasis.

### Adipocyte GPD2 deficiency impairs energy expenditure

To investigate the role of GPD2 in adipocytes, we generated adipocyte-specific GPD2 knockout (GPD2^AKO^) mice by crossing GPD2^flox/flox^ mice with adiponectin-Cre mice. *Gpd2* expression was selectively ablated in adipose tissues, with no detectable changes in other tissues (Fig. S1A). When fed a normal chow diet (NCD), GPD2^AKO^ mice exhibited body weights comparable to those of WT littermates (Fig. S1B), and no significant differences were observed in the mass of the three major adipose depots (Fig. S1C). To evaluate adaptive thermogenesis, we performed cold tolerance tests. Following acclimation at thermoneutrality (30°C), acute cold exposure resulted in significantly lower rectal and surface temperatures in GPD2^AKO^ mice compared with WT controls (Fig. 2A and B). Consistent with this impaired thermogenic capacity, indirect calorimetry showed that GPD2^AKO^ mice exhibited reduced oxygen consumption (VO_2_), CO_2_ production (VCO_2_), and heat production under both room temperature and thermoneutral conditions (Fig. 2C-E). Notably, these metabolic defects in GPD2^AKO^ mice were markedly accentuated upon cold stimulation (Fig. 2C-E). GPD2^AKO^ mice exhibited increased BAT mass (Fig. 2F). Compared with WT controls, the BAT of knockout mice appeared enlarged and whitened, reflecting excessive lipid accumulation (Fig. 2G-H and Fig. S1D). Consistent with this morphological alteration, the oxygen consumption rate of BAT was dramatically reduced in GPD2^AKO^ mice (Fig. 2I), collectively indicating impaired BAT function. In contrast, neither the mass nor the morphology of iWAT or eWAT showed significant changes (Fig. 2F-H and Fig. S1D). Given that GPD2 is a mitochondrial protein, we next investigated the impact of its ablation on mitochondrial structure and function. Transmission electron microscopy of BAT showed that, while mitochondrial abundance remained comparable between genotypes, cristae density was notably reduced and mitochondrial ultrastructure displayed swelling in GPD2^AKO^ mice (Fig. 2J and K). Moreover, upon acute cold exposure, UCP1 expression in BAT was significantly repressed in GPD2-deficient mice, as well as cristae assembly-related proteins including CHCHD3 and CHCHD6 (Fig. 2L and Fig. S1E).

**Fig. 2.**
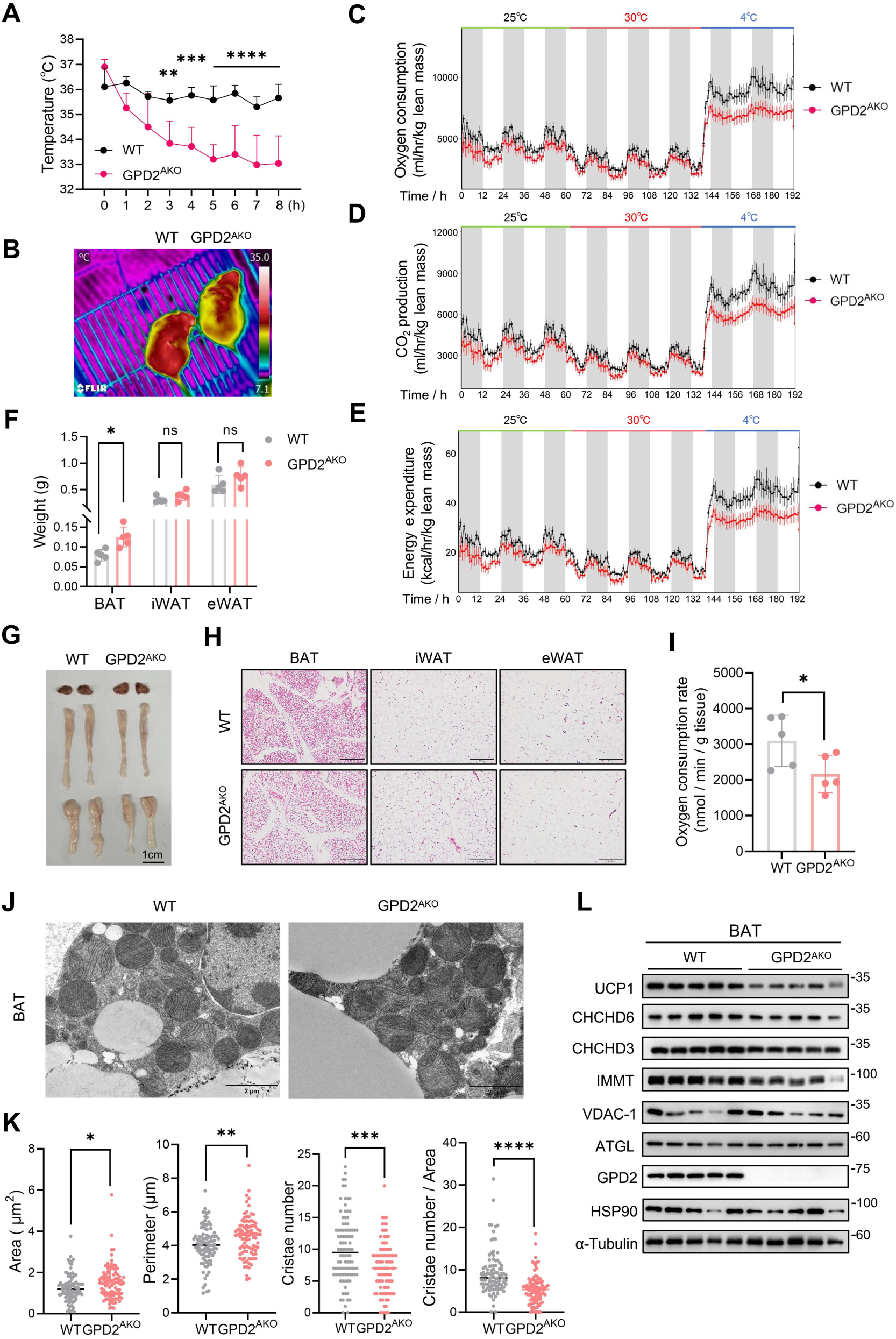
GPD2^AKO^ mice exhibit impaired energy expenditure and thermogenesis. Twelve-week-old male WT and GPD2^AKO^ mice were housed under TN conditions (30°C) for two weeks, then exposed to 4°C, and subsequently subjected to the following measurements. (A) Rectal temperature measurements under 4°C exposure (n = 5). Data were analyzed using two-way ANOVA with Bonferroni’s multiple comparisons test. (B) Representative infrared thermal images of the dorsal surface of the mice after 8 h of exposure to 4°C. (C-E) Oxygen consumption (C), CO₂ production (D), and energy expenditure (E) in 12-week-old male WT and GPD2^AKO^ mice (n = 6/group) under room temperature, TN, and cold exposure conditions. (F-G) Tissue weights of BAT, iWAT, and eWAT (F) and representative images of adipose tissues (G). Data in (F) were analyzed using two-tailed unpaired Student’s t-test (BAT and eWAT) or Mann-Whitney test (iWAT). (H) Representative H&E staining images of iWAT, eWAT, and BAT (scale bar, 200 μm). (I) OCR of BAT from male WT and GPD2^AKO^ mice (n = 5/group) after 8 h of exposure to 4°C. Data were analyzed using two-tailed unpaired Student’s t-test. (J) Mitochondrial ultrastructure in BAT of WT and GPD2^AKO^ mice under cold exposure after TN acclimation. Scale bar, 2 μm. (K) Quantitative analysis of mitochondrial area, perimeter, cristae number, and cristae density from the transmission electron microscopy images. Data were analyzed using two-tailed unpaired Student’s t-test (perimeter) or Mann-Whitney test (area, cristae number, cristae density). (L) Western blot analysis of UCP1 and the mitochondrial cristae structural components in BAT from mice under cold exposure after TN acclimation. *, p < 0.05; **, p < 0.01; ***, p < 0.001.

Given that the acute cold challenge primarily reflects BAT function, we next asked whether GPD2 disruption also impairs WAT browning in response to chronic cold exposure. Following a 3-day cold challenge, GPD2^AKO^ mice exhibited increased iWAT mass and enlarged lipid droplets compared with WT controls (Fig. S2A-C). In line with the observations in BAT, a similar reduction in UCP1 levels was observed in iWAT of GPD2^AKO^ mice upon prolonged cold stimulation, as indicated by the UCP1 immunohistochemical staining and western blot analysis (Fig. S2B and D). These data together suggested that GPD2 deficiency also attenuated the browning process of WAT.

Having observed defective adaptive thermogenesis in GPD2^AKO^ mice, we next assessed whether this impairment was cell-autonomous. To address this, we manipulated GPD2 expression in an *in vitro* adipocyte model. Neither overexpression nor knockout of GPD2 affected the adipogenic differentiation (Fig. S3A and B), indicating that GPD2 was dispensable for adipocyte differentiation. Notably, Seahorse analysis showed that cellular OCR was markedly reduced in GPD2-deficient adipocytes (Fig. 3A). Measurement of mitochondrial respiration in mature adipocytes using substrates respectively fed into complex Ⅰ, complex Ⅱ, and fatty acid β-oxidation revealed that GPD2 disruption significantly attenuated oxygen consumption in isolated mitochondria (Fig. 3B-D). Furthermore, ectopic expression of GPD2 in C3H10T1/2 cells increased UCP1 levels (Fig. 3E and F). Conversely, disruption of GPD2, achieved through either siRNA-mediated transient knockdown or sgRNA-mediated stable knockout, consistently reduced UCP1 expression in immortalized brown adipocytes (Fig. 3G-I). Supporting this, GPD2 expression positively correlated with UCP1 levels in mouse BAT (Fig. 3J). Collectively, these findings established that GPD2 cell-autonomously maintained thermogenic capacity and energy homeostasis in adipocytes.

**Fig. 3.**
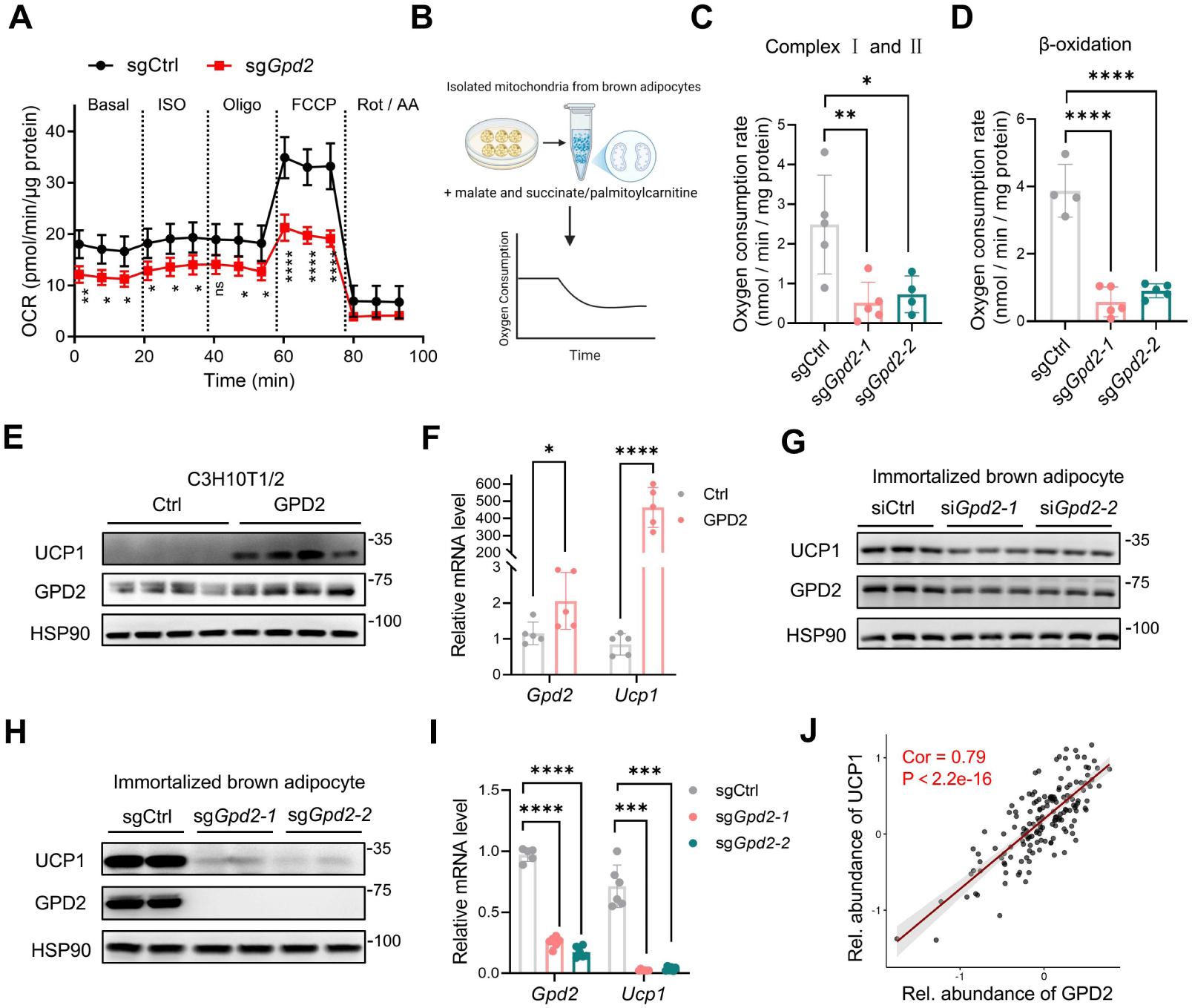
GPD2 cell-autonomously maintains thermogenic capacity and energy homeostasis. (A) Seahorse analysis of OCR in immortalized brown adipocytes from sgCtrl and sg*Gpd2-1* groups (n = 5-6). Data were analyzed using two-way ANOVA with Bonferroni’s multiple comparisons test. (B-D) Mitochondria were isolated from immortalized brown adipocytes of sgCtrl and sg*Gpd2* groups following isoproterenol treatment for 16 h. OCR for complex I/II (C) and fatty acid oxidation (D) were measured (n = 4-5 per group). Data were analyzed by ordinary one-way ANOVA with Bonferroni’s multiple comparisons test. (E-F) C3H10T1/2 cells overexpressing GPD2 and control cells were differentiated into adipocytes. On day 8 post-adipogenic differentiation, UCP1 and GPD2 expressions were analyzed by western blot (E) and qPCR (F) (n=5) after treatment with isoproterenol for 16 h. Data in (F) were analyzed using two-tailed unpaired Student’s t-test. (G) Immortalized brown preadipocytes were transfected with siNC or si*Gpd2* on day 3 of differentiation and treated with isoproterenol on day 6 for 16 h. UCP1 and GPD2 protein expression were assessed. (H-I) UCP1 and GPD2 expression were analyzed by western blot (H) and qPCR (I) in immortalized brown adipocytes from sgCtrl and sg*Gpd2* groups following isoproterenol treatment for 16 h (n=5-6). Data in (I) were analyzed by Brown–Forsythe and Welch ANOVA tests followed by Tamhane’s T2 multiple comparisons test. (J) Correlation between UCP1 and GPD2 protein levels in mouse BAT from the OPABAT correlation network (n = 163 mice). *, p < 0.05; **, p < 0.01; ****, p < 0.0001.

### Loss of GPD2 inhibits glycolysis in BAT

To explore the mechanisms underlying thermogenic dysfunction in GPD2^AKO^ mice, we performed RNA sequencing (RNA-seq) on BAT samples. Transcriptomic analysis identified 1,570 upregulated and 1,365 downregulated differentially expressed genes (DEGs) in GPD2^AKO^ mice compared with WT (Fig. S4A). Remarkably, both KEGG pathway analysis and Gene Set Enrichment Analysis (GSEA) revealed significant downregulation of pathways related to thermogenesis, oxidative phosphorylation, pyruvate metabolism, citrate cycle (TCA cycle), and glycerolipid metabolism (Fig. 4A and Fig. S4B-F). Genes involved in these pathways, including *Ucp1, Cidea, Cox6c, Ndufb11, Sdha, Uqcrc2, Dlat, Cs, Cox4i1, Dgat1,* and *Agpat2,* were downregulated upon GPD2 ablation (Fig. 4B). These transcriptomic changes were validated by qPCR analysis (Fig. 4C). We then performed targeted metabolomics on BAT from WT and GPD2^AKO^ mice to further investigate the metabolic alteration of GPD2 deficiency. As expected from its role in catalyzing the conversion of G3P to DHAP^13^, GPD2 ablation resulted in G3P accumulation (Fig. 4D), thereby confirming the reliability of our metabolomic profiling. Notably, the analysis revealed a widespread reduction in glycolytic and TCA cycle intermediates following GPD2 ablation, including fructose 6-phosphate, pyruvate, lactate, acetyl-CoA, fumarate, and oxaloacetate (Fig. 4D). To assess whether the reduced metabolite levels reflect impaired glycolytic function, we measured extracellular acidification rate (ECAR) and confirmed that GPD2 deficiency significantly reduced glycolytic capacity, as indicated by the diminished increase in ECAR upon glucose and oligomycin treatment (Fig. 4E and F).

**Fig. 4.**
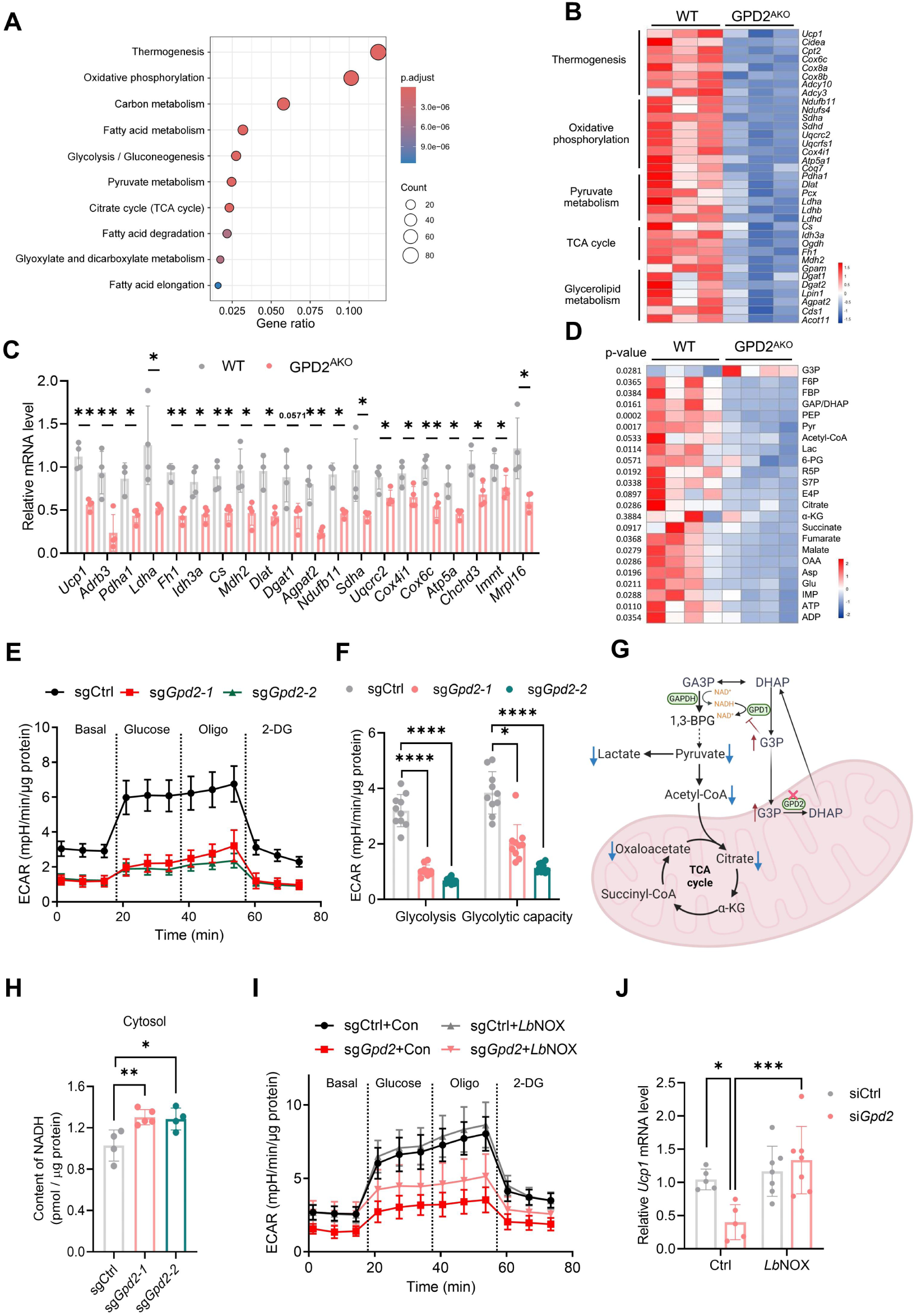
GPD2 regulates glucose oxidative metabolism. (A) Significantly downregulated KEGG pathways among DEGs in BAT of GPD2^AKO^ mice after 8 h of exposure to 4°C, following two-week TN acclimation. (B) Heatmap of downregulated genes in BAT from male GPD2^AKO^ mice compared with WT controls (n = 3/group). (C) qPCR validation of altered genes in BAT of GPD2^AKO^ mice (n=3-4). Data were analyzed using two-tailed unpaired Student’s t-test (*Ucp1*, *Adrb3*, *Idh3a*, *Cs*, *Mdh2*, *Agpat2*, *Uqcrc2*, *Cox4i1*, *Cox6c*, *Chchd3*, *Immt*, and *Mrpl16*), Mann-Whitney test (*Dgat1*, *Sdha*) or Welch’s correction (*Pdha1*, *Ldha*, *Fh1*, *Dlat*, *Ndufb11*, and *Atp5a*). (D) Metabolomics analysis of BAT from 12-week-old WT and GPD2^AKO^ mice (n = 4). (E) ECAR measured by Serhorse analyzer in immortalized brown adipocytes from sg-Ctrl and sg-*Gpd2* groups (n = 10). (F) Glycolysis and glycolytic capacity were calculated based on ECAR measurements under basal and stimulated conditions (n = 10). Data of glycolysis were analyzed by Brown–Forsythe and Welch ANOVA tests followed by Tamhane’s T2 multiple comparisons test. Data of glycolytic capacity were analyzed by Kruskal–Wallis test followed by Dunn’s multiple comparisons test. (G) Schematic representation of the glycerol-3-phosphate shuttle. GPD2 deficiency leads to G3P accumulation in the cytosol. Elevated G3Ps, the product of GPD1, inhibited GPD1 activity and promoted NADH levels. Increased NADH suppressed glycolysis and reduced the metabolites. (H) Cytosolic NADH levels in immortalized brown adipocytes from sg-Ctrl and sg-*Gpd2* groups (n = 4-5). Data were analyzed using ordinary one-way ANOVA followed by Bonferroni’s multiple comparisons test. (I) ECAR in brown adipocytes from sgCtrl and sg*Gpd2-1* groups, with or without *Lb*NOX overexpression (n = 9-10). (J) qPCR analysis of *Ucp1* mRNA levels in *Lb*NOX-overexpressing immortalized brown adipocytes with *Gpd2* deficiency (si*Gpd2-1*), compared to control group (n = 5-7). Data were analyzed using ordinary one-way ANOVA followed by Bonferroni’s multiple comparisons test. *, p < 0.05; **, p < 0.01; ***, p < 0.001; ****, p < 0.0001.

To elucidate how GPD2 disruption inhibits glycolysis, we measured cytosolic NADH levels, given their critical role in glycolytic flux and their potential modulation by GPD2^13, 18^. GPD2 deficiency led to a significant increase in G3P, which is the product of GPD1, inhibited GPD1 activity and in turn elevated the cytosolic NADH content (Fig. 4G and H). Considering that cytosolic NADH inhibits GAPDH, this increase in NADH likely accounts for the observed glycolytic suppression. To test whether NADH accumulation is indeed causally responsible for glycolytic defects, we expressed *Lactobacillus brevis* NADH oxidase (*Lb*NOX), which oxidizes NADH to NAD^+ 23^, in GPD2-deficient brown adipocytes. *Lb*NOX overexpression effectively reduced the cytosolic NADH levels in GPD2-KO cells, without affecting mitochondrial NADH (Fig. S4G-I). Remarkably, *Lb*NOX expression rescued the impaired ECAR and totally reversed the reduction in *Ucp1* expression caused by GPD2 deficiency (Fig. 4I and J). These data collectively demonstrated that GPD2 ablation suppresses glucose catabolism through NADH-mediated glycolytic inhibition.

### GPD2 shapes the epigenetic landscape to control thermogenic gene expression

Given that GPD2 disruption reduces glycolytic and TCA cycle metabolites, many of which, including lactate, acetyl-CoA, and fumarate, are established modulators of histone modifications, we hypothesized that GPD2 remodels the epigenetic landscape to control thermogenic gene expression. Initial western blot analysis of BAT revealed an obvious decrease in the levels of H3K27ac upon GPD2 deficiency, whereas the levels of histone H3 lysine 18 lactylation (H3K18la), histone H3 lysine 4 monomethylation (H3K4me1), and histone H3 lysine 27 trimethylation (H3K27me3) remained unchanged (Fig. 5A). H3K27ac is a marker of active promoters and enhancers. To investigate the genome-wide changes in this activating histone mark, we performed Cleavage Under Targets and Tagmentation (CUT&Tag) analysis of H3K27ac on BAT from WT and GPD2^AKO^ mice. Loss of GPD2 led to a reduction in H3K27ac occupancy at numerous genomic loci (Fig. 5B). Consistently, ATAC-seq revealed decreased chromatin accessibility at numerous genomic regions in GPD2^AKO^ mice (Fig. 5C). Differential peak analysis of H3K27ac CUT&Tag and ATAC-seq data showed that these downregulated peaks were predominantly enriched in intergenic regions (Fig. S5A). To link these distal regulatory elements to their potential target genes, we annotated the differential peaks to their nearest genes. We then integrated H3K27ac CUT&Tag, ATAC-seq, and RNA-seq data. This integrative analysis revealed that genes co-downregulated across all three omics datasets were functionally enriched for thermogenesis and oxidative phosphorylation (Fig. 5D and S5B), including *Ucp1, Cox6c, Sdhd, Coq7,* and *Mrpl35*.

**Fig. 5.**
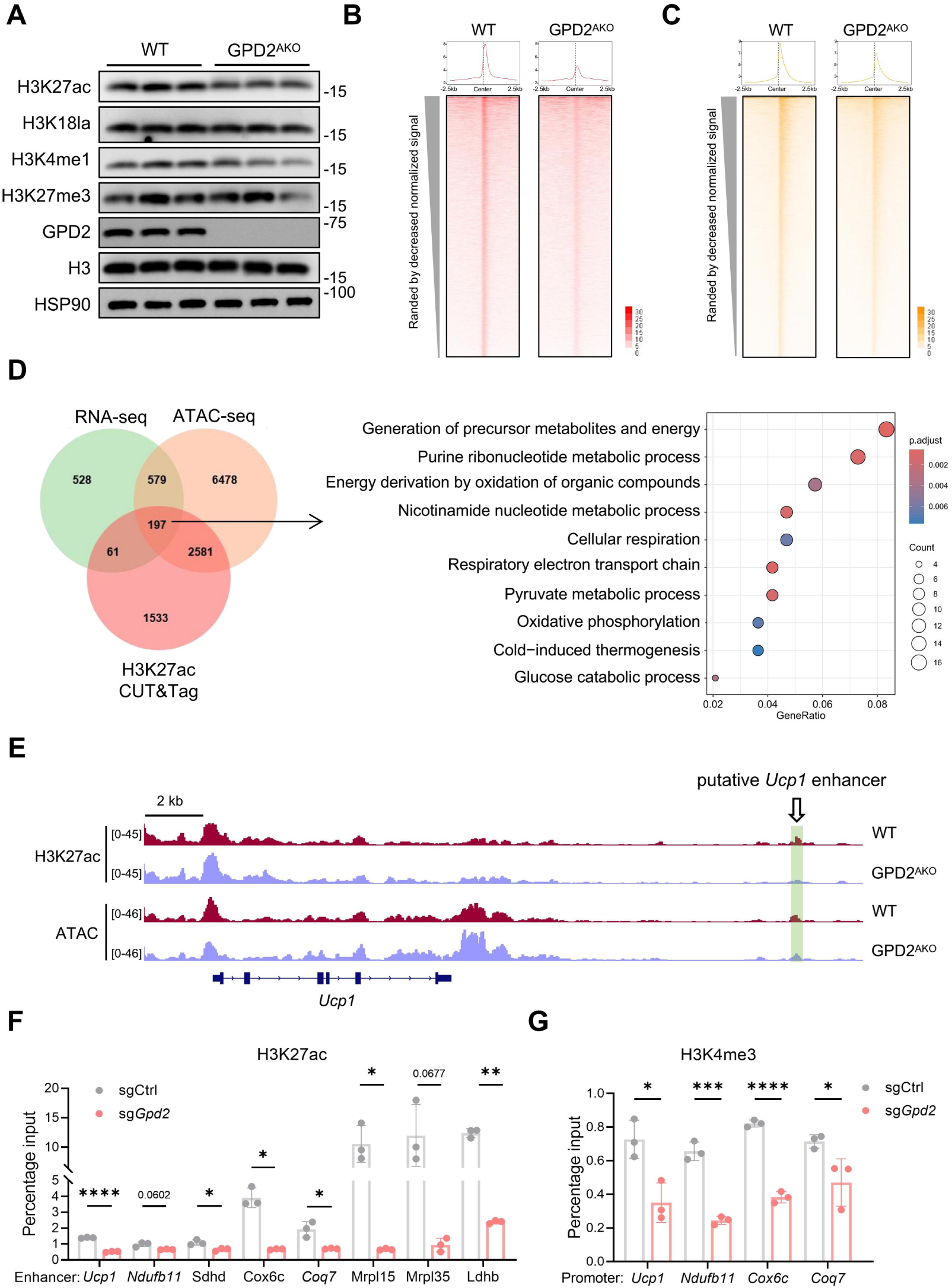
GPD2 remodels the epigenetic landscape of BAT. (A) Western blot analysis of common histone modifications in BAT from male GPD2^AKO^ and WT mice (n = 3) after 8 h cold exposure, following two-week TN acclimation. Histone H3 and HSP90 served as loading controls. (B-C) Metaplots and heatmap showing signals of downregulated regions for H3K27ac and ATAC in the BAT from WT and GPD2^AKO^ mice. (D) Venn diagrams showing co-downregulated genes across RNA-seq, H3K27ac CUT&Tag (annotated to nearest genes), and ATAC-seq (annotated to nearest genes) in BAT of GPD2^AKO^ versus WT mice. Dot plot showing GO pathway enrichment analysis of the overlapping gene set. (E) H3K27ac and ATAC signals at *Ucp1* enhancer loci in BAT from WT and GPD2^AKO^ mice. The putative enhancer regions were indicated by green rectangles. The merged display of two biological replicates was shown. (F-G) ChIP-qPCR validation of H3K27ac (F) and H3K4me3 (G) enrichment at indicated regions in immortalized brown adipocytes from control and sg-*Gpd2-1* groups treated with isoproterenol for 16 h (n = 3). Data in (F) were analyzed using two-tailed unpaired Student’s t-test (*Ucp1*, *Ndufb11*, *Sdhd*) or Welch’s correction (*Cox6c*, *Coq7*, *Mrpl15*, *Mrpl35*, *Ldhb*). Data in (G) were analyzed using two-tailed unpaired Student’s t-test. *, p < 0.05; **, p < 0.01; ****, p < 0.0001.

In the H3K27ac CUT&Tag analysis, we identified a putative *Ucp1* enhancer region (chr8:83,309,946−83,310,225) located 19.7 kb downstream of *Ucp1* gene locus. This region exhibited H3K27ac enrichment in WT BAT, which was markedly reduced upon GPD2 deficiency (Fig. 5E). Importantly, ATAC-seq confirmed open chromatin at the same site (Fig. 5E). Consistent with this, public databases further revealed that chromatin accessibility at this locus was increased in both iWAT and BAT following cold stimulation (Fig. S6A)^24^, correlating with the increased expression of UCP1. Additionally, published ChIP-seq datasets also showed H3K4me1 enrichment at this locus in BAT, another enhancer-associated mark, which was further augmented by cold exposure (Fig. S6A)^25^. The presence of active histone marks H3K27ac and H3K4me1 together with high chromatin accessibility collectively indicated an active enhancer at this genomic region. We then performed ChIP-qPCR to validate the decrease in H3K27ac at the putative enhancer regions of *Ucp1* by GPD2 disruption, as well as other thermogenesis-related genes (Fig. 5F). Consistent with the silencing of enhancer activity, ChIP-qPCR also revealed reduced H3K4me3 occupancy at the promoter regions of *Ucp1* (Fig. 5G).

To directly test whether the putative enhancer region possesses intrinsic transcriptional activity, we cloned this fragment into a pGL3-Promoter luciferase reporter vector driven by an SV40 basal promoter. The resulting construct was transiently transfected into brown adipocytes, and luciferase activity was measured to assess enhancer function. Compared with the empty vector (EV) control, the fragment containing the putative enhancer significantly increased luciferase activity (Fig. 6A), indicating that this region possesses enhancer activity capable of driving transcription in a heterologous promoter context. We next sought to functionally validate the enhancer activity of this region for *Ucp1* expression. To this end, we designed sgRNAs (sgeh1 and sgeh2) targeting this locus and used CRISPR-dCas9-LSD1 and CRISPR-dCas9-KRAB-MeCP2 systems to repress its enhancer function in brown adipocytes (Fig. S6B). The dCas9-LSD1 system was employed to remove enhancer-associated active histone marks, and dCas9-KRAB-MeCP2 system was used to induce heterochromatin formation, which indeed reduced H3K27ac enrichment at the *Ucp1* putative enhancer region (Fig. S6C). Strikingly, both epigenetic editing approaches that suppressed this locus markedly inhibited UCP1 expression, at both mRNA and protein levels (Fig. 6B-E). To complement these loss-of-function data, we used the CRISPR-dCas9-p300 system to enhance H3K27ac at this potential enhancer region and observed a significant increase in UCP1 expression (Fig. S6D-G), collectively confirming that this region functions as a bona fide enhancer for *Ucp1*. More importantly, targeted restoration of H3K27ac at this enhancer region significantly rescued the reduction in UCP1 expression caused by GPD2 knockdown (Fig. 6F and G). Consistently, replenishing the acetyl-CoA pool with sodium acetate (NaAc) treatment also reversed the decreased UCP1 levels in GPD2-deficient adipocytes (Fig. 6H-J). Taken together, these data demonstrated that GPD2 shaped the epigenetic landscape to control thermogenic gene expression.

**Fig. 6.**
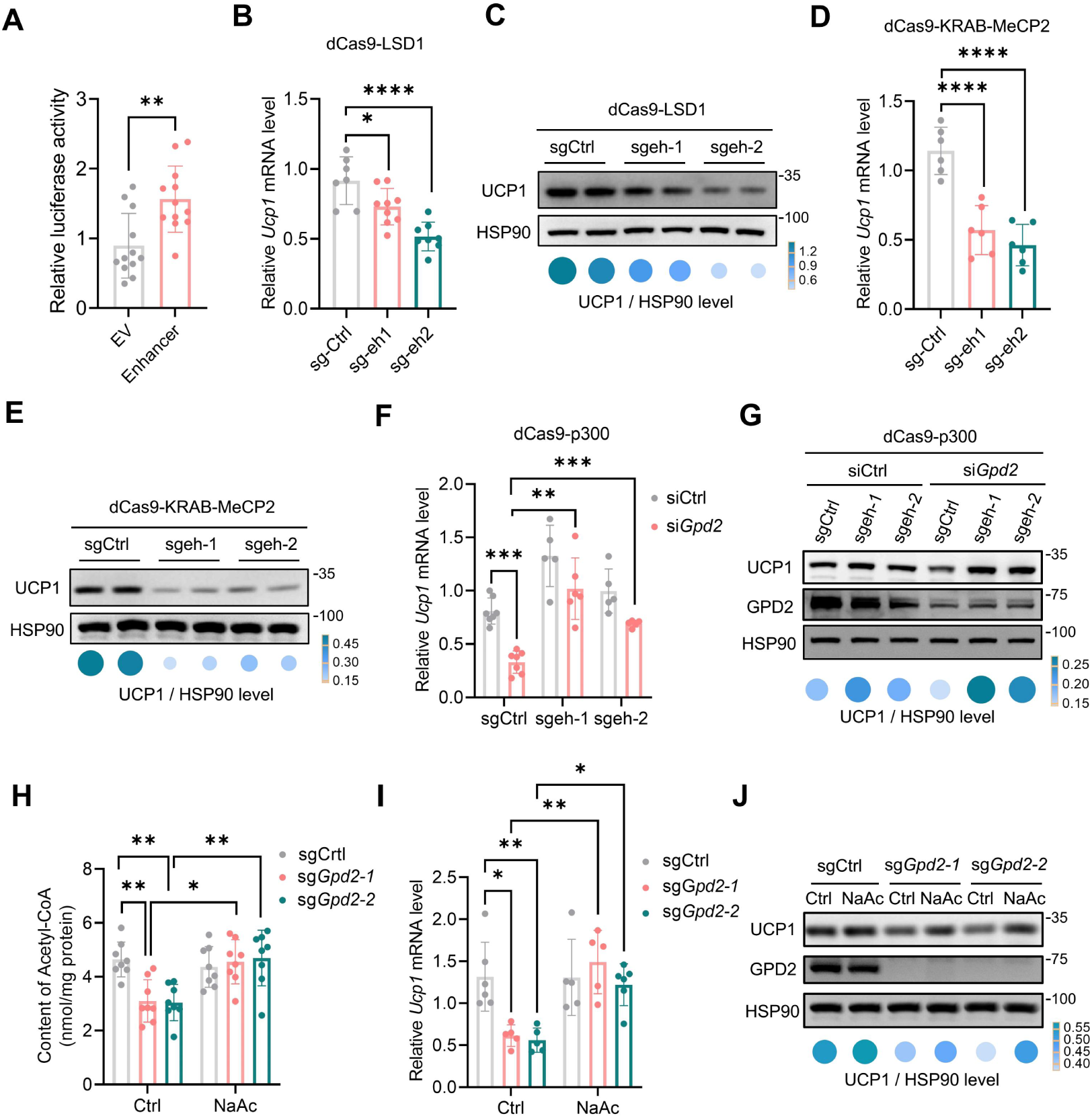
Epigenetic editing of the enhancer validates GPD2-dependent UCP1 regulation. (A) Immortalized brown preadipocytes were transfected with either the pGL3-Promoter control EV or the same vector containing a putative enhancer fragment, followed by luciferase reporter assay (n=12). Data were analyzed by the Mann–Whitney test. (B-C) Repression of putative enhancer loci in immortalized brown adipocytes using the dCas9-LSD1 system. The mRNA (B, n = 7-9) and protein (C) levels of UCP1 were assessed. Data in (B) were analyzed using ordinary one-way ANOVA followed by Bonferroni’s multiple comparisons test. (D-E) Repression of putative enhancer loci in immortalized brown adipocytes using the dCas9-KRAB-MeCP2 system. UCP1 mRNA (D, n = 6) and protein (E) levels were assessed. Data in (D) were analyzed using ordinary one-way ANOVA followed by Bonferroni’s multiple comparisons test. (F-G) The dCas9-p300 system was used to restore H3K27ac at putative enhancer loci in GPD2-deficient brown adipocytes (si*Gpd2-1*). UCP1 mRNA and protein levels were then measured by qPCR (F, n = 5-7) and western blot (G), respectively. Data in (F) were analyzed using ordinary one-way ANOVA followed by Bonferroni’s multiple comparisons test. (H) Acetyl-CoA levels in isoproterenol-treated immortalized brown adipocytes from sg-Ctrl and sg-*Gpd2* groups, with or without sodium acetate treatment (n = 8). Data were analyzed by ordinary one-way ANOVA followed by Bonferroni’s multiple comparisons test. (I-J) qPCR (I, n=5-7) and western blot (J) and analysis of UCP1 in GPD2-deficient cells with or without sodium acetate treatment. Data in (I) were analyzed by ordinary one-way ANOVA followed by Bonferroni’s multiple comparisons test. *, p < 0.05; **, p < 0.01; ***, p < 0.001.

### GPD2^AKO^ mice are prone to developing metabolic disorders

Having established that GPD2 regulated thermogenic activation through a metabolic-epigenetic axis, we next examined the metabolic phenotype of GPD2^AKO^ mice under high-fat diet (HFD) challenge. Upon HFD feeding, GPD2^AKO^ male mice exhibited significantly higher body weights than WT controls, accompanied by increased BAT mass and pronounced lipid accumulation (Fig. 7A-C). Metabolic cage analysis showed that GPD2^AKO^ mice had lower whole-body O_2_ consumption, CO_2_ production, and energy expenditure than WT mice (Fig. 7D-F). Consistently, ex vivo oxygen consumption assays revealed that even at room temperature, BAT from GPD2^AKO^ mice displayed significantly reduced respiratory activity compared with that from WT controls (Fig. 7G), collectively indicating functional deterioration of BAT. Although glucose tolerance was not markedly impaired in GPD2^AKO^ mice, their insulin sensitivity was obviously reduced (Fig. 7H and I). Further serum biochemical analysis showed that triglycerides (TG), total cholesterol (TC), high-density lipoprotein (HDL), and low-density lipoprotein (LDL) levels were significantly higher in GPD2^AKO^ mice than controls, whereas blood glucose and liver injury markers (ALT and AST) remained comparable (Fig. 7J-N). Notably, these metabolic abnormalities in male GPD2^AKO^ mice were consistently recapitulated in female mice (Fig. S7A-C). Collectively, these findings demonstrated that GPD2 deficiency exacerbated BAT dysfunction under HFD conditions, leading to more severe metabolic disturbances, including obesity, dyslipidemia, and insulin resistance.

**Fig. 7.**
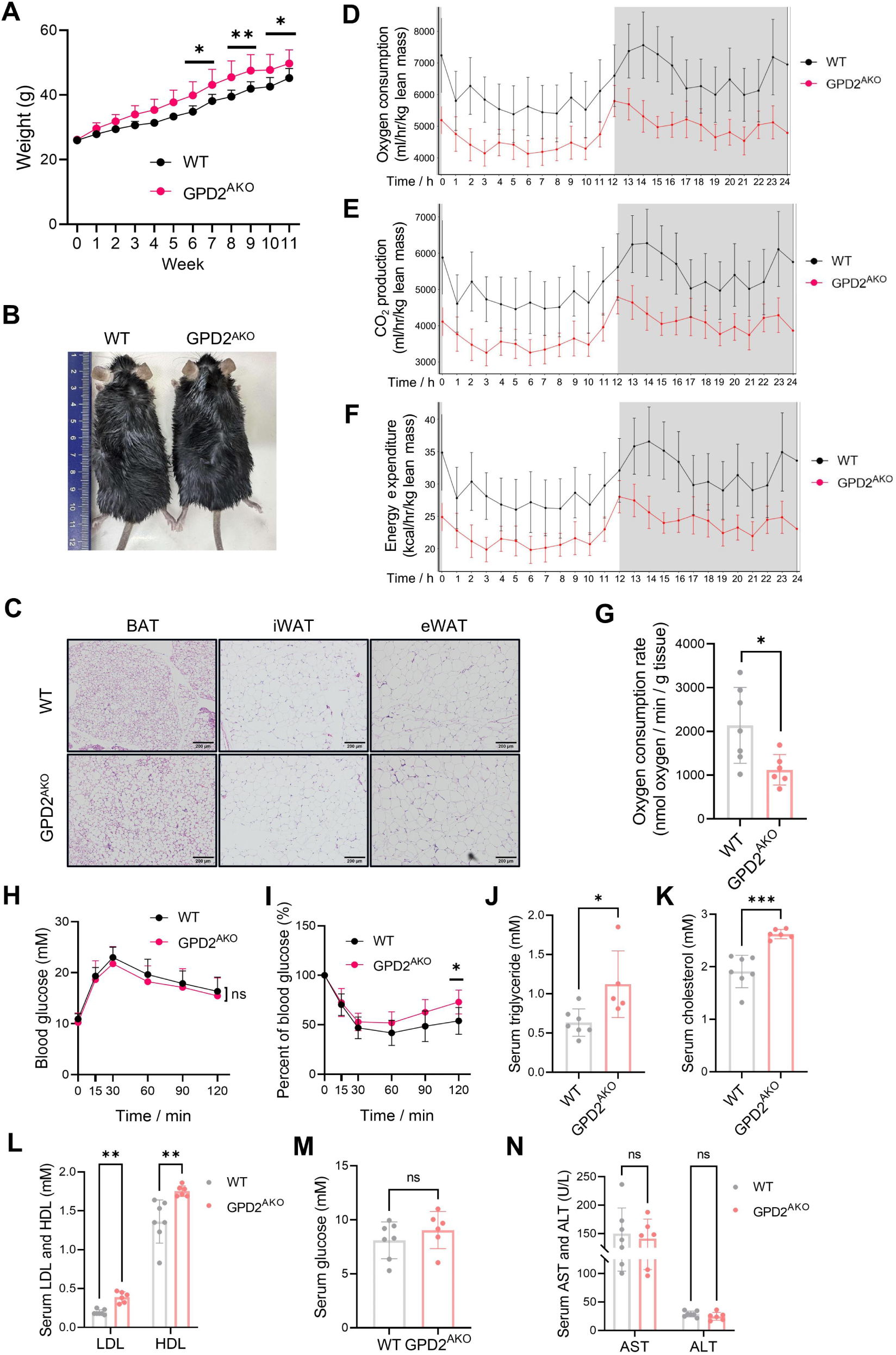
GPD2^AKO^ mice exhibit increased susceptibility to metabolic disorders. (A-B) WT and GPD2^AKO^ mice were fed an HFD starting at 8 weeks of age (n = 5-7). Body weight changes (A) and representative whole-body photographs (B) were shown. Data in (A) were analyzed using two-way ANOVA followed by Bonferroni’s multiple comparisons test. (C) Representative H&E staining of iWAT, eWAT, and BAT in WT and GPD2^AKO^ mice fed with HFD (scale bar, 200 μm). (D-F) Oxygen consumption (D), CO_2_ production (E), and energy expenditure (F) were assessed at room temperature in WT and GPD2^AKO^ mice (n = 6/group). (G) OCR in BAT from HFD-fed WT and GPD2^AKO^ mice (n = 6-7). Data were analyzed using two-tailed unpaired Student’s t-test. (H-I) GTT (H) and ITT (I) in WT and GPD2^AKO^ mice fed with HFD (n = 6-8). Data were analyzed using two-way ANOVA followed by Bonferroni’s multiple comparisons test. (J-N) Serum levels of triglycerides (J), total cholesterol (K), LDL and HDL (L), glucose (M), AST, and ALT (N) in male WT and GPD2^AKO^ mice. Data were analyzed using two-tailed unpaired Student’s t-test, Mann-Whitney test (LDL) or Welch’s correction (serum cholesterol and glucose). *, p < 0.05; **, p < 0.01.

## Discussion

The mammalian NADH shuttle system comprises two primary pathways: the G3PS and MAS. Beyond its canonical role in transferring cytosolic reducing equivalents, the G3PS serves as a metabolic hub that integrates glycolysis, lipid synthesis, and oxidative phosphorylation. Unlike the MAS, the G3PS transfers electrons from NADH to FAD, producing FADH_2_ and one less ATP per NADH. This bioenergetic inefficiency may dissipate energy as heat, representing a UCP1-independent thermogenic mechanism. Alongside this intrinsic property, we demonstrate that GPD2, the core component of the G3PS, controls thermogenic activation by remodeling the epigenetic landscape in brown adipocytes. GPD2 governs metabolite availability to drive H3K27ac deposition and UCP1 expression. Thus, the G3PS couples energy inefficiency with GPD2-mediated epigenetic control to coordinate both UCP1-independent and UCP1-dependent thermogenesis. In addition to G3PS, it has been reported that glutamic-oxaloacetic transaminase (GOT1), a key MAS enzyme, is induced by cold in BAT. Moreover, the upregulation of GOT1 in BAT activates the MAS, thereby facilitating mitochondrial fatty acid oxidation while simultaneously suppressing glucose oxidation. GOT1-mediated substrate preference in BAT consequently enhances thermogenesis^26,27^. Together with our current findings, these studies highlight the profound role of NADH shuttle system in coupling substrate flux with energy expenditure.

Previous studies have unveiled the physiopathological significance of GPD2. In diabetic kidney disease, GPD2 deficiency exacerbates podocyte injury through mechanisms involving mitochondrial dysfunction and oxidative stress^17^. Similarly, in the liver, loss of GPD2 contributes to hepatic steatosis, highlighting its role in systemic lipid homeostasis^15^. Furthermore, GPD2 modulates macrophage inflammatory responses, underscoring its function in immune regulation^18^. Our current study broadens the understanding of GPD2 function by defining a previously unrecognized, cell-autonomous role in thermogenic fat activation. Glucose fuels thermogenesis through glycolysis and subsequent mitochondrial oxidation, which is essential both for providing reducing equivalents for uncoupled respiration and for generating metabolites that actively regulate thermogenic programming^28, 29^. We demonstrate that in thermogenic adipocytes, GPD2 governs a metabolic-epigenetic axis by regulating glycolytic flux. GPD2 deficiency leads to NADH accumulation in the cytosol, which suppresses glycolysis and the availability of acetyl-CoA, thus inhibiting thermogenic gene expression via epigenetic reprogramming (Fig. S8). This finding emphasizes that intracellular metabolite flux functions as a direct and rapid epigenetic regulator in thermogenic activation. The functional importance of acetyl-CoA was further demonstrated by NaAc supplementation. Replenishing the intracellular acetyl-CoA pool successfully reversed the downregulation of thermogenic genes caused by GPD2 deficiency. Consistent with this model, we confirmed that modulating histone acetylation at the potential enhancer regions of *Ucp1* directly affects its expression, therefore identifying a previously undefined enhancer locus for this gene. These results provide direct causal evidence that the availability of key metabolites, particularly acetyl-CoA, is mechanistically essential for sustaining the thermogenic program in adipocytes.

GPD2 catalyzes the conversion of G3P to DHAP. Accordingly, metabolomic profiling in the present study revealed marked G3P accumulation upon GPD2 knockout. This observation is further supported by human genetic data. Specifically, a serum metabolome GWAS of 6,136 Finnish males identified a GPD2 SNP (rs201060271) as the strongest genome-wide association for circulating G3P levels^30^. Emerging evidence positions G3P beyond its conventional role as a biosynthetic precursor. G3P can be utilized for glycerolipid synthesis, converted to DHAP to enter glycolysis, or dephosphorylated to glycerol to support gluconeogenesis^31^. The obesity and hyperlipidemia phenotype observed in GPD2^AKO^ mice may thus arise, at least in part, from the effect of accumulated G3P on these pathways, including glycerolipid synthesis and gluconeogenesis. Beyond its role as a biosynthetic precursor, G3P also acts as a signaling molecule. Recent studies have identified G3P as a specific ligand for the carbohydrate response element-binding protein (ChREBP), directly linking its accumulation to the induction of hepatic *de novo* lipogenesis^32^. Moreover, elevated G3P levels have been identified not only as biomarkers of cellular senescence but also as active inducers of the senescent state^33, 34^. Given its dual role as a metabolic hub and signaling metabolite, a key unanswered question is whether G3P accumulation directly regulates thermogenic capacity, potentially acting as a metabolic signal that modulates energy expenditure in adipose tissue.

In summary, we elucidate a critical role for GPD2 in regulating thermogenic fat activation. GPD2 governs glucose oxidative process and the production of acetyl-CoA, which in turn shape the histone modification landscape to activate core thermogenic gene expression. Loss of GPD2 impairs thermogenic capacity and leads to systemic metabolic dysfunction. This study establishes a direct mechanistic link between glycolytic metabolism and epigenetic control of adipose thermogenesis, revealing GPD2 as a fundamental node integrating fuel utilization with transcriptional regulation. Collectively, our findings open new avenues for improving metabolic homeostasis by targeting GPD2-mediated metabolite-epigenome axis in adipose tissues.

## Methods

### Animal model

GPD2^flox/flox^ mice were kindly provided by Dr. Tiffany Horng (ShanghaiTech University). GPD2^AKO^ mice were generated by crossing homozygous GPD2^flox/flox^ mice with adiponectin-Cre mice. WT C57BL/6J mice were purchased from GemPharmatech Co., Ltd. (Nanjing, China). All animal experiments were approved by the Fudan University Shanghai Medical College Animal Care and Use Committee (No. 20240408-001) and followed the National Institutes of Health guidelines on the care and use of animals. Mice were housed under a 12-h light/dark cycle with free access to food and water. Mice in the room temperature group were housed in ventilated cages at 21±1°C and a humidity of 50%±5%, whereas those in thermoneutral group were maintained at 30°C.

### Cell culture and treatment

The basal culture medium consisted of DMEM (Meilunbio, MA0212) or DMEM/F-12 (for primary cells; meilunbio, MA0214), supplemented with 10% fetal bovine serum (Gibco, 10091148) and 1% penicillin/streptomycin (Meilunbio, MA0110). Immortalized brown preadipocytes were established and differentiated as previously described^35, 36^. The day when cells reached 80% confluence was designated as day -2. Cells were then cultured in a differentiation medium consisting of the basal culture medium supplemented with 1 μg/mL insulin and 1 nM triiodothyronine. On day 0, the medium was replaced with an induction cocktail: differentiation medium further supplemented with 0.5 mM isobutylmethylxanthine and 0.5 μM dexamethasone. Cells were maintained in differentiation medium on days 2 and 4 and considered mature on day 6 for subsequent analyses. For C3H10T1/2 adipogenic induction, day 0 was designated two days after contact inhibition. Differentiation was initiated using the same induction cocktail as described above. Cells were maintained in the basal differentiation medium on days 2, 4, and 6, and considered mature on day 8 for subsequent treatments. For isolation of the SVF from BAT or iWAT, tissues were minced and digested at 37°C for 40-50 min in serum-free DMEM containing 0.075% collagenase (Sigma, C2139). The digest was filtered through a cell strainer and centrifuged. The resulting pellet was resuspended, plated, and washed the following day to remove non-adherent cells. The adherent, proliferating cells were considered adipose-derived stem cells and were induced to differentiate using protocols identical to those described above for brown and white preadipocytes cell lines.

### Manipulation of GPD2 expression in cell models

To generate GPD2 knockout cells, we employed the CRISPR-Cas9 system. Complementary oligonucleotides targeting GPD2 were phosphorylated and annealed using T4 PNK enzyme (#M0201S, New England Biolabs). The resulting double-stranded DNA product was ligated into the BsmBI-digested lentiCRISPR v2 vector using T4 DNA ligase (#M0202S, New England Biolabs). The sequences of the complementary oligonucleotides for each target gene are detailed in Supplementary Table S1. For GPD2 knockdown experiments, transfections were performed on day 3 of differentiation. GPD2-specific siRNA oligonucleotides and negative control were synthesized by Shanghai GenePharma Co., Ltd. with the sequence listed in Supplementary Table S1. For overexpression experiments, the full-length *Gpd2* coding sequence was inserted into the PCDH-CMV-MCS-EF1-CoPGFP-T2A-Puro vector. For lentivirus production, the lentiviral plasmid was co-transfected with packaging vectors into HEK293T cells. Viral supernatants were collected at 48 and 96 h post-transfection, filtered through a 0.45 μm filter, and used to infect immortalized preadipocytes for 48 hours. Stable knockout or overexpression cells were selected with puromycin (10 µg/ml).

### CRISPR interference (CRISPRi) and CRISPR activation (CRISPRa) *in vitro*

For CRISPRi and CRISPRa in cell-based experiments, lentiviruses encoding dCas9-KRAB-MeCP2, dCas9-LSD1, or dCas9-p300 were transduced into immortalized pre-brown adipocytes. Transduced cells were selected with 5 µg/mL blasticidin to establish stable cell lines expressing the respective dCas9 constructs. The lentiGuide-Puro vector, containing sgRNAs targeting the *Ucp1* enhancer regions, was then transduced into these stable lines, followed by puromycin selection. The sgRNA sequences are listed in Supplementary Table S1.

### Mitochondrial isolation

BAT or cultured cells were homogenized in mitochondrial isolation buffer containing 300 mM sucrose, 10 mM HEPES, and 0.2 mM EDTA (pH 7.2). The homogenate was centrifuged at 600 × g for 10 min at 4°C to remove tissue and cellular debris. The resulting supernatant was then centrifuged at 8,000 × g for 10 min at 4°C to pellet the mitochondrial fraction. The supernatant after this step was collected as the crude cytosolic fraction for subsequent measurements. The mitochondrial pellet was washed and resuspended in an appropriate buffer for subsequent analyses.

### Tissue and mitochondrial oxygen consumption assay

To measure tissue respiration, adipose tissue fragments were suspended in oxygen-saturated assay buffer consisting of PBS supplemented with 2% bovine serum albumin (BSA), 25 mM glucose, and 1 mM pyruvate. Oxygen consumption rates were then measured using a Clark-type oxygen electrode (Oxygraph+ system, Hansatech). For isolated mitochondria, respiratory activity was assessed in PBS buffer containing 2% BSA supplemented with 0.5 mM malate and 10 mM succinate. To assess β-oxidation capacity, isolated mitochondria were stimulated with 0.5 mM malate and 50 µM palmitoylcarnitine. Oxygen consumption rates were normalized to tissue weight or to the protein content of cells or mitochondria, as appropriate for each experimental condition.

### Seahorse assay

Cellular respiration and glycolytic capacity in brown adipocytes were measured using a Seahorse XFe96 Analyzer (Agilent). Immortalized brown adipocytes at day 4 of adipogenic differentiation were trypsinized, seeded into Seahorse XF 96-well microplates at equal cell numbers per well, and cultured for an additional 48 h under differentiation conditions prior to the assay. For extracellular acidification rate (ECAR) measurements, cells were equilibrated in serum-free Seahorse XF medium containing 2 mM glutamine at 37 °C for 60 min. ECAR was measured with Seahorse XF Glycolysis Stress Test Kit (Agilent) and determined under basal conditions and sequentially following the addition of glucose (10 mM), oligomycin (1 µM), and 2-deoxy-D-glucose (2-DG, 50 mM), according to the manufacturer’s instructions. Basal glycolysis was calculated as the ECAR following glucose injection minus the basal ECAR. Glycolytic capacity was calculated as the maximal ECAR value following oligomycin injection minus the basal ECAR. For OCR measurements, the assay was performed in Seahorse XF medium supplemented with 10 mM glucose, 2 mM glutamine, and 1 mM pyruvate. OCR was measured under basal conditions and sequentially following the injection of isoproterenol (10 µM), oligomycin (0.5 µM), carbonylcyanide-p-trifluoromethoxyphenylhydrazone (FCCP, 0.5 µM), and a mixture of rotenone and antimycin A (0.5 µM). Both the ECAR and OCR were normalized to the protein levels.

### Cold tolerance assay

Mice acclimated to thermoneutral conditions (30℃) for 14 days were subjected to acute cold exposure at 4°C. Rectal temperature was measured hourly using a rectal probe connected to an electronic thermometer (Alcott Biotech, Shanghai, China). Body surface temperature was monitored and recorded using an infrared digital thermography camera (T430SC, Teledyne FLIR).

### Indirect calorimetric assessment

Oxygen consumption (VO₂), carbon dioxide production (VCO₂), and heat production were measured using a Comprehensive Lab Animal Monitoring System (Columbus Instruments), as previously described^22, 37^. Mice were individually housed in the recording chambers and allowed to acclimate for 24 h prior to data collection. Measurements were performed under a 12 h light/dark cycle with ad libitum access to food and water. The light phase was from 7:00 a.m. to 7:00 p.m., and the dark phase was from 7:00 p.m. to 7:00 a.m. next day. VO₂, VCO₂, and heat production were all normalized to lean body mass.

### Chromatin Immunoprecipitation (ChIP)

Cells were cross-linked by incubation with 1% formaldehyde in PBS for 10 min at room temperature. The reaction was quenched by adding glycine to a final concentration of 125 mM for 5 min. Cells were then lysed in ChIP lysis buffer, and chromatin was sonicated using a M220 Focused Ultrasonicator (Covaris) to shear DNA into fragments of 200-1000 bp. After removal of cell debris, an aliquot of the supernatant was saved as the input control. The remaining lysate was incubated overnight at 4°C with rotation in the presence of the target-specific primary antibody. The following day, immune complexes were captured by incubation with Protein A/G agarose beads. The beads were then washed five times with ChIP wash buffer. Cross-links were reversed by incubation at 70°C overnight. Samples were then treated sequentially with RNase A and Proteinase K. DNA was purified using MiniElute PCR purification kits (Qiagen) and quantified. Subsequently, 1 ng of the ChIP-enriched DNA and corresponding input DNA were analyzed by qPCR.

### Glucose tolerance test (GTT), insulin tolerance test (ITT), and serum biochemical analysis

GTT and ITT were performed as previously described^38^. For GTT, mice were fasted for 16 h (from 5:00 p.m. to 9:00 a.m. the following day) and then received an intraperitoneal injection of D-glucose at a dose of 2 mg/g body weight for mice on NCD or 1.5 mg/g body weight for mice fed an HFD. Blood glucose levels were measured using a glucometer (Roche) via tail vein sampling at 0, 15, 30, 60, 90, and 120 min after glucose administration. For ITT, mice were fasted for 4 h (from 10:00 a.m. to 2:00 p.m.) and then received an intraperitoneal injection of recombinant human insulin (Novo Nordisk) at a dose of 0.75 IU/kg body weight. Blood glucose levels were measured at the same time intervals as in GTT. For serum biochemical analysis, levels of blood glucose, TG, TC, HDL, and LDL were measured using an automated biochemistry analysis device (Roche) according to the manufacturer’s instructions.

### Transmission electron microscopy and quantification

BAT was carefully dissected and cut into small pieces (approximately 1 mm^3^) with minimal mechanical damage. Tissue samples were fixed in electron microscopy fixative containing 2.5% glutaraldehyde for 2 h at room temperature, followed by overnight incubation at 4°C. Subsequent sample processing, sectioning, and imaging were performed by Wuhan Servicebio Technology Co., Ltd. Mitochondrial ultrastructure was analyzed using ImageJ software. Mitochondrial area was measured by delineating the outer membrane contour using the freehand selection tool. Mitochondrial cristae density was calculated as the number of cristae per unit mitochondrial area.

### RNA sequencing

Total RNA was extracted from BAT of WT and GPD2^AKO^ mice using TRIzol reagent. RNA quantity and integrity were assessed using the RNA Nano 6000 Assay Kit on an Agilent 2100 Bioanalyzer system (Agilent Technologies, CA, USA). High-quality RNA was used for subsequent library preparation. Library construction, sequencing, and bioinformatic analysis were performed by Metabo-Profile Biotechnology Co., Ltd. (Shanghai, China).

### Metabolomics analysis

Freshly isolated BAT was snap-frozen in liquid nitrogen and homogenized in ice-cold 80% methanol (HPLC grade) at 10 mL per gram of tissue. The homogenate was incubated overnight at −80°C. The following day, the sample was centrifuged at 12,000 × g for 20 min at 4°C. The resulting supernatant was collected, lyophilized, and reconstituted in acetonitrile. Subsequent detection and analysis were performed by the Core Facility of Shanghai Medical College, Fudan University.

### CUT&Tag and ATAC-seq

Nuclei were freshly isolated from BAT using tissue nuclei isolation kit (12514ES56, Yeasen) and used for library preparation with the Hyperactive Universal CUT&Tag Assay Kit for Illumina Pro (TD904, Vazyme). Briefly, Concanavalin A (ConA) beads were activated using cold bead activation buffer. Isolated nuclei were incubated with the activated ConA beads and then resuspended in pre-chilled antibody buffer. For each sample, 1 μg of primary antibody against the target antigen was added, and the mixture was incubated overnight at 4°C with rotation. The following day, a secondary antibody was added to capture the primary antibody-chromatin complexes. Subsequently, pA/G-Tnp Pro was added and incubated for 1 hour at room temperature to facilitate binding. Chromatin fragmentation was carried out by incubation at 37°C for 1 h. The resulting fragmented DNA was purified, and libraries were constructed and amplified using the TruePrep Index Kit V2 for Illumina (TD202, Vazyme). Library preparation was performed using less than 30 ng of fragmented DNA. Final libraries were quantified using Qubit and the Agilent Tapestation system. Sequencing was subsequently performed by Novogene Co., Ltd. (Beijing, China).

ATAC-Seq was performed using the Hyperactive ATAC-Seq Library Prep Kit (TD711-01, Vazyme). Briefly, nuclei isolated from BAT were resuspended in 50 µL of transposition reaction mix. The transposition reaction was incubated at 37°C for 30 min, followed by DNA extraction, library construction, and purification. Library quality was assessed using a BioAnalyzer 2100 system (Agilent Technologies, Santa Clara, CA) prior to deep sequencing. Sequencing was performed by Novogene Co., Ltd. (Beijing, China).

For bioinformatics analysis, raw CUT&Tag and ATAC-seq reads were filtered using fastp (version 0.23.2)^39^ and then mapped to the reference genome (mm10) using Bowtie2 (version 2.3.5.1)^40^, as previously described^41, 42^. Aligned reads were filtered for a minimum MAPQ of 30 using SAMtools (version 1.3.1)^43^. MarkDuplicates tool in Picard was used to identify and remove PCR duplicates from the aligned reads. MACS2 was applied on each bam file to call peaks with q-value threshold 0.05 ^44^. The BigWig files were generated by deepTools (version 3.5.1) using RPKM option of bamCoverage command^45^. The differential peaks were defined based on the following criteria: for ATAC-seq, regions with a fold change > 1.2 between GPD2^AKO^ and WT samples; for H3K27ac CUT&Tag, regions with a fold change > 1.5. Identification of the nearest genes to differential peaks was performed using the annotatePeaks.pl script in HOMER^46^.

### RNA extraction and real-time qPCR

Total RNA was extracted from tissues or cultured cells using TRIzol reagent. RNA was reverse-transcribed into complementary DNA using a HiScript III 1st Strand cDNA Synthesis Kit (Vazyme, R312-01) according to the manufacturer’s instructions. Quantitative real-time PCR was performed on a QuantStudio^TM^ real-time PCR system using SYBR Green Supermix (Vazyme, Q711-02). Relative gene expression was analyzed using the 2^−ΔΔCT^ method, with 18S rRNA as the endogenous control. Expression levels were normalized to the control group. Primer sequences are listed in Table S2.

### Western blot

Tissues or cultured cells were lysed in lysis buffer containing 2% SDS and 50 mM Tris-HCl (pH 6.8), supplemented with a protease inhibitor cocktail (Roche, Cat# 4693159001). Protein concentrations of the lysates were determined using BCA protein assay kit. Equal amounts of protein were separated by SDS-PAGE and subsequently transferred to polyvinylidene difluoride (PVDF) membranes (Millipore Corp., Bedford, MA). Membranes were blocked with 5% (w/v) skimmed milk in TBST for 2 hours at room temperature, followed by overnight incubation at 4°C with primary antibodies. After washing with TBST, membranes were incubated with the corresponding horseradish peroxidase (HRP)-conjugated secondary antibodies (Jackson ImmunoResearch) for 1 hour at room temperature. Protein bands were visualized using an enhanced chemiluminescence (ECL) detection kit (ShareBio, Cat# SB-WB012) and imaged with an ImageQuant LAS 4000 imaging system (GE Healthcare Life Sciences). The antibodies used in the study were listed in Table S3.

### H&E staining and immunohistochemistry

Histological analyses were performed by Shanghai Ruiyu Biotechnology Co., Ltd. Briefly, adipose tissues were dissected and immediately fixed in 4% paraformaldehyde solution for 24 h. Fixed tissues were embedded in paraffin, sectioned, and subjected to hematoxylin and eosin (H&E) staining or immunohistochemical staining according to standard protocols. Quantitative analysis of adipocyte diameter in adipose tissues was performed using ImageJ software.

### Statistical Analysis

All statistical analyses were performed using GraphPad Prism (GraphPad Software, San Diego, CA). Data are presented as mean ± SD. Comparisons between two groups were analyzed using two-tailed unpaired Student’s t-test for normally distributed data with equal variances, or two-tailed unpaired Student’s t-test with Welch’s correction for data with unequal variances. For non-normally distributed data, the Mann–Whitney U test was used. For comparisons involving more than two groups, one-way ANOVA followed by Bonferroni’s post hoc test was applied for normally distributed data with equal variances. When variances were unequal, the Brown–Forsythe and Welch ANOVA tests followed by Tamhane’s T2 multiple comparisons test were used. For non-normally distributed data, the Kruskal–Wallis test followed by Dunn’s multiple comparisons test was performed. For experiments with two independent variables, two-way ANOVA followed by Bonferroni’s multiple comparisons test was used. In all analyses, p<0.05 was considered statistically significant. The specific statistical tests used for each figure panel are indicated in the corresponding figure legends.

## Author contributions

Y.L. and Q.-q.T. initiated and supervised the project; Y.L., Y.-j.S., and M.D. designed the research; Y.-j.S., M.D., L.-x.X., X.D., H.-y.X., Y.-f.X., Y.-f.K., Z.-y.M., J.-y.X., L.-y.Z., W.-j.S., S.-w.Q., Y.T. performed the experiments; G.Z. carried out bioinformatics analysis. Y.-j.S., M.D., G.Z. and Y.L. analyzed data; and Y.L. and Y.-j.S. wrote the manuscript.

## Acknowledgments

We are grateful to Dr. Tiffany Horng from ShanghaiTech University for providing GPD2^flox/flox^ mice. This work was supported by National Natural Science Foundation of China (NSFC) Grants 82522019 (to Y. L.), Noncommunicable Chronic Diseases-National Science and Technology Major Project 2026ZD0556400 (to Y. L.), NSFC Grants 82370881 (to Y. L.), 92457301 and 82330023 (to Q.-Q. T.), 82300959 (to X.D.), and High-Level Local University Development Funding.

---

**Fig. S1.**
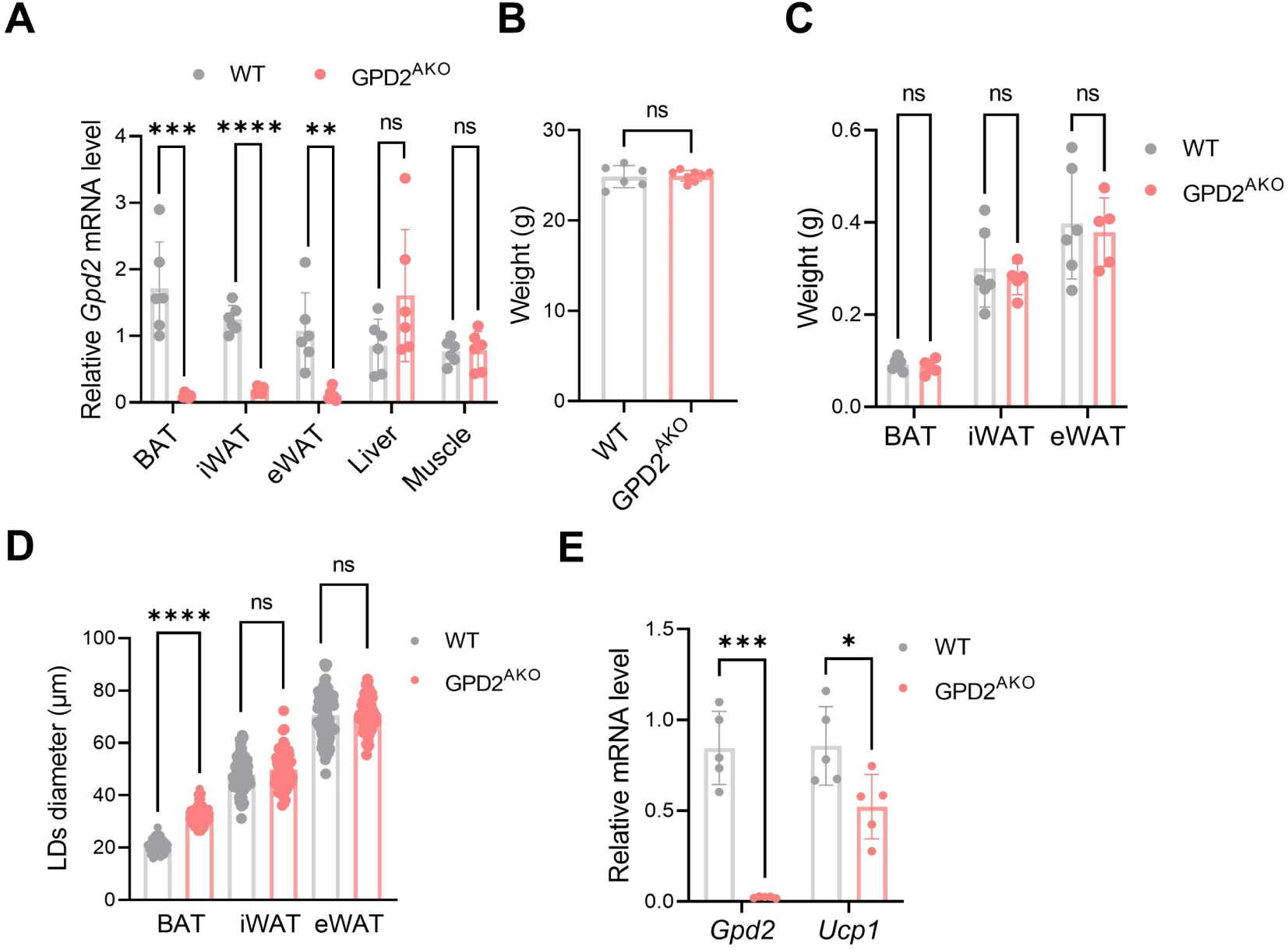
GPD2^AKO^ mice exhibit BAT degeneration. (A) *Gpd2* mRNA levels in various tissues of 12-week-old WT and GPD2^AKO^ mice (n = 6). Data were analyzed using two-tailed unpaired Student’s t-test (liver and muscle) or Welch’s correction (BAT, iWAT, and eWAT). (B-C) Body weight (B, n = 6-8) and tissue weights of BAT, iWAT, and eWAT (C, n = 5-6) in WT and GPD2^AKO^ mice housed at room temperature under NCD. Data in (B) and (C) were analyzed using two-tailed unpaired Student’s t-test. (D) Quantification of lipid droplet diameters from H&E staining of adipose tissues using ImageJ software. Each dot represents an individual lipid droplet. Data were analyzed using two-tailed unpaired Student’s t-test with Mann-Whitney test. (E) qPCR analysis of *Gpd2* and *Ucp1* in BAT from male WT and GPD2^AKO^ mice (n = 5/group) under cold exposure after TN acclimation. Data were analyzed using two-tailed unpaired Student’s t-test (*Ucp1*) or Welch’s correction (*Gpd2*). *, p < 0.05; **, p < 0.01; ***, p < 0.001; ****, p < 0.0001.

**Fig. S2.**
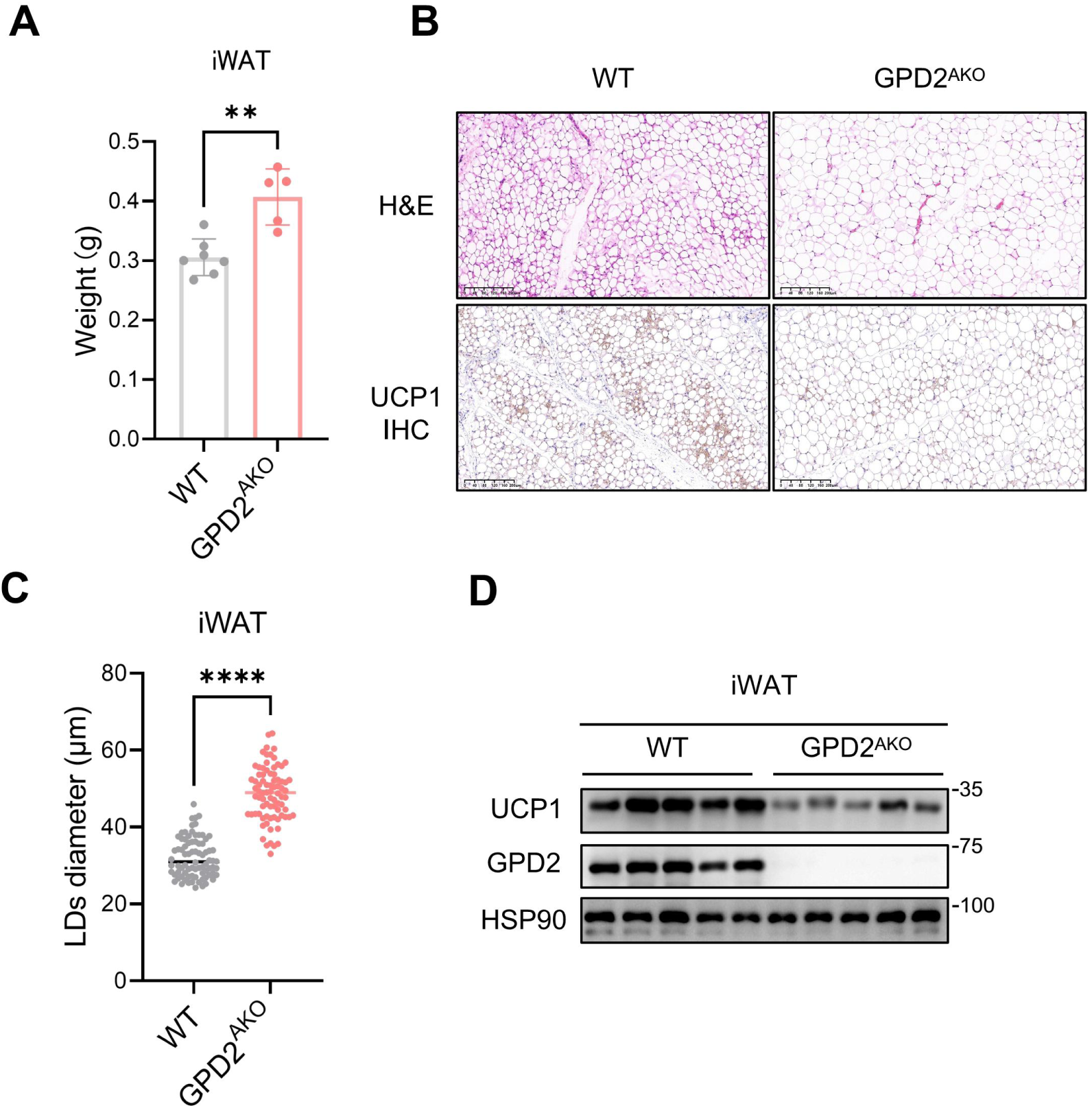
GPD2 deficiency suppresses the browning capacity of WAT. (A) iWAT weight in 12-week-old WT and GPD2^AKO^ mice following 3 days of cold exposure (n = 5-7). Data were analyzed using two-tailed unpaired Student’s t-test. (B-C) Representative images of H&E staining and UCP1 IHC in iWAT of WT and GPD2^AKO^ mice following 3 days of cold exposure (B) (scale bar, 200 μm). Quantification of lipid droplet diameter from H&E staining (C). Data in (C) were analyzed using two-tailed unpaired Student’s t-test. (D) Western blot analysis of UCP1 and GPD2 in iWAT (n = 5). **, p < 0.01; ****, p < 0.0001.

**Fig. S3.**
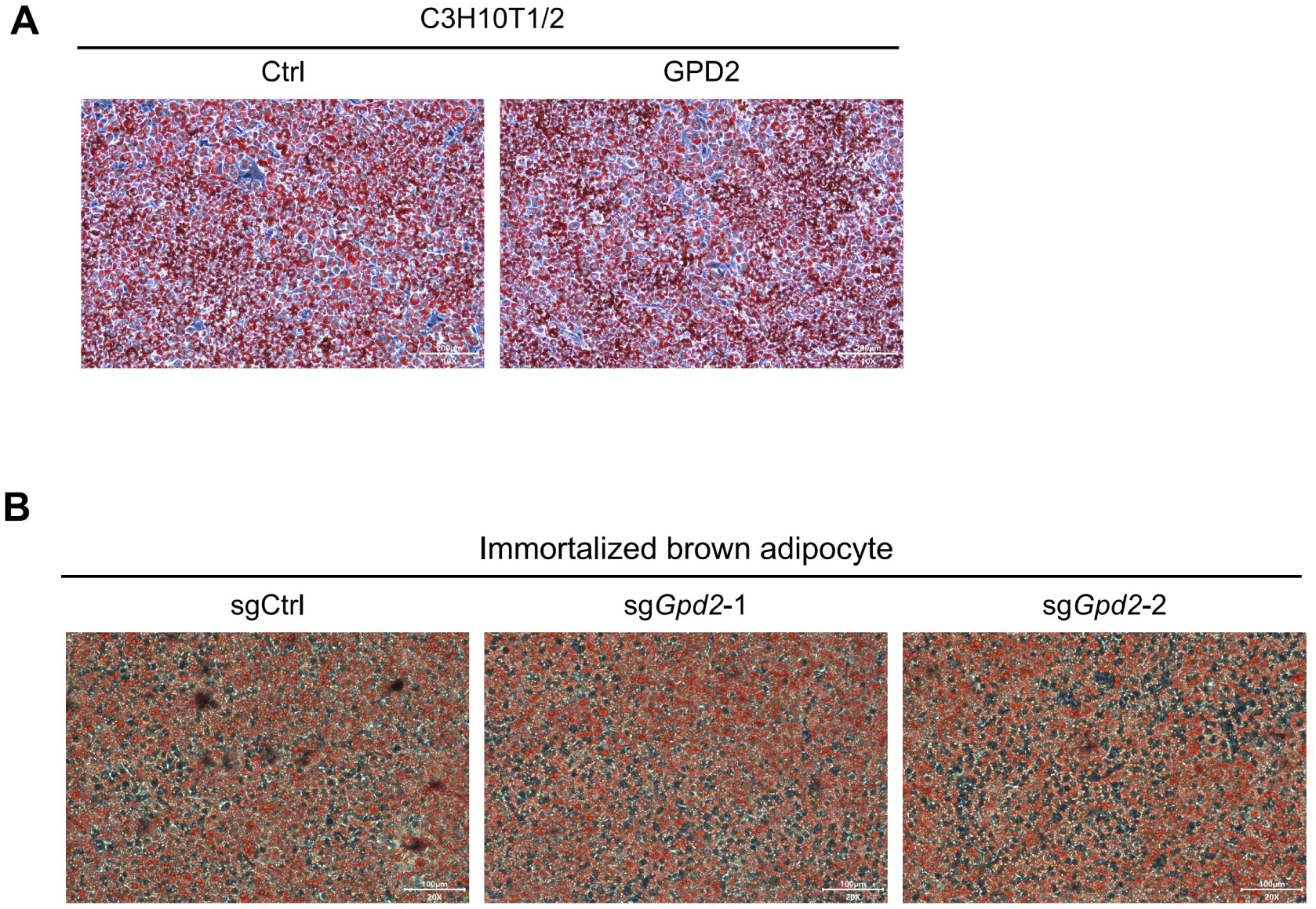
GPD2 has no effect on adipogenic differentiation. (A) Oil Red O staining of C3H10T1/2 cells following adipogenic differentiation upon GPD2 overexpression (scale bar, 200 μm). (B) Oil Red O staining of immortalized brown adipocytes following adipogenic differentiation upon GPD2 knockout (scale bar, 100 μm).

**Fig. S4.**
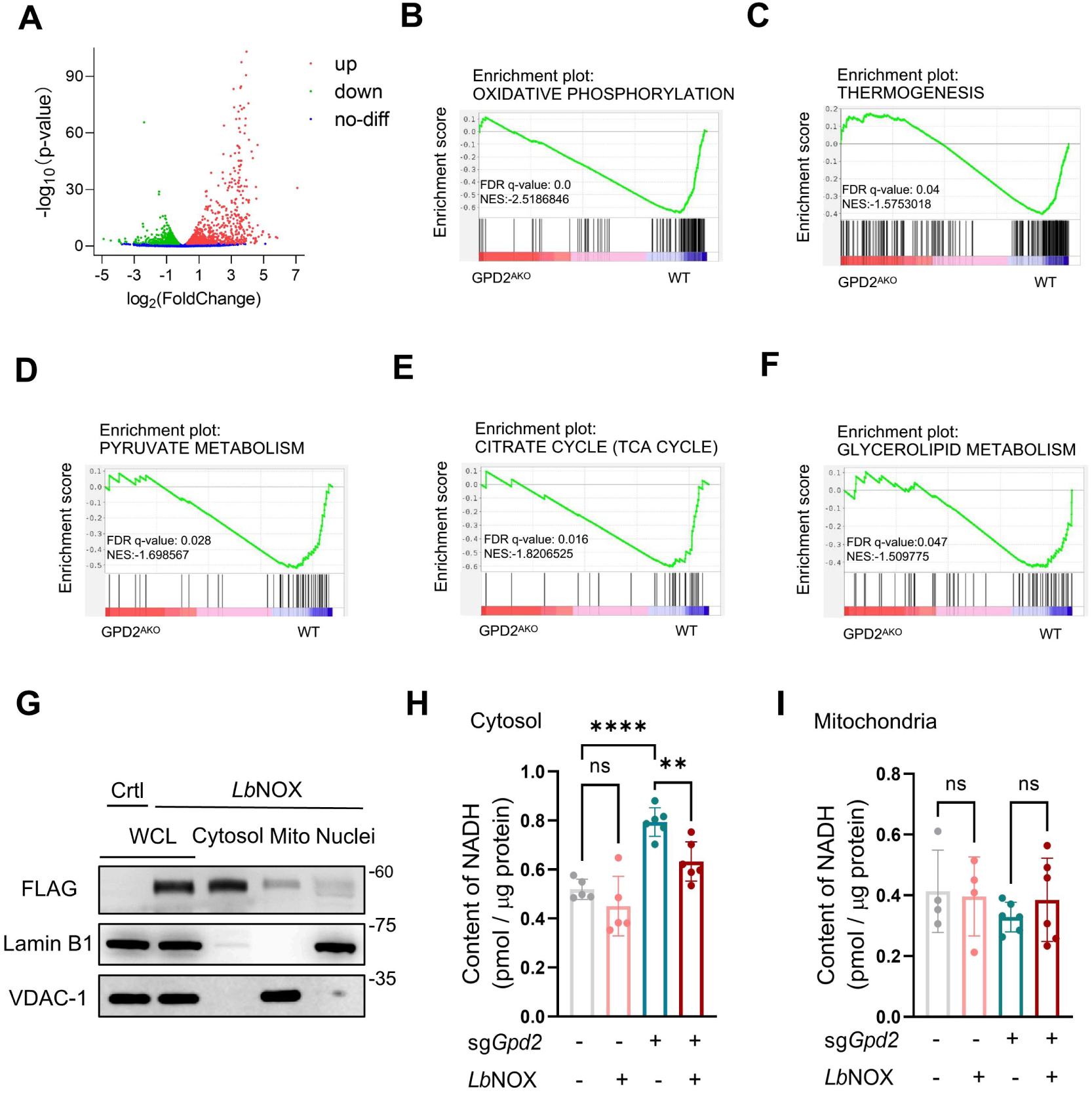
Key pathways related to energy metabolism are inhibited by GPD2 disruption. (A) Volcano plot showing DEGs between GPD2^AKO^ and WT mice. A total of 1,570 upregulated and 1,365 downregulated genes were identified (P < 0.05). (B-F) GSEA showing enrichment of various pathways in BAT of GPD2^AKO^ mice compared with WT. (G) Western blot analysis of *Lb*NOX-FLAG protein levels in whole cell lysates (WCL), cytosol, mitochondria, and nuclei from control and *Lb*NOX-overexpressing immortalized brown adipocytes. (H-I) Cytosolic (H) and mitochondrial (I) NADH levels in sgCtrl and sg*Gpd2-1* brown adipocytes with or without *Lb*NOX overexpression (n = 4-6). Data were analyzed using ordinary one-way ANOVA followed by Bonferroni’s multiple comparisons test. **, p < 0.01; ****, p < 0.0001.

**Fig. S5.**
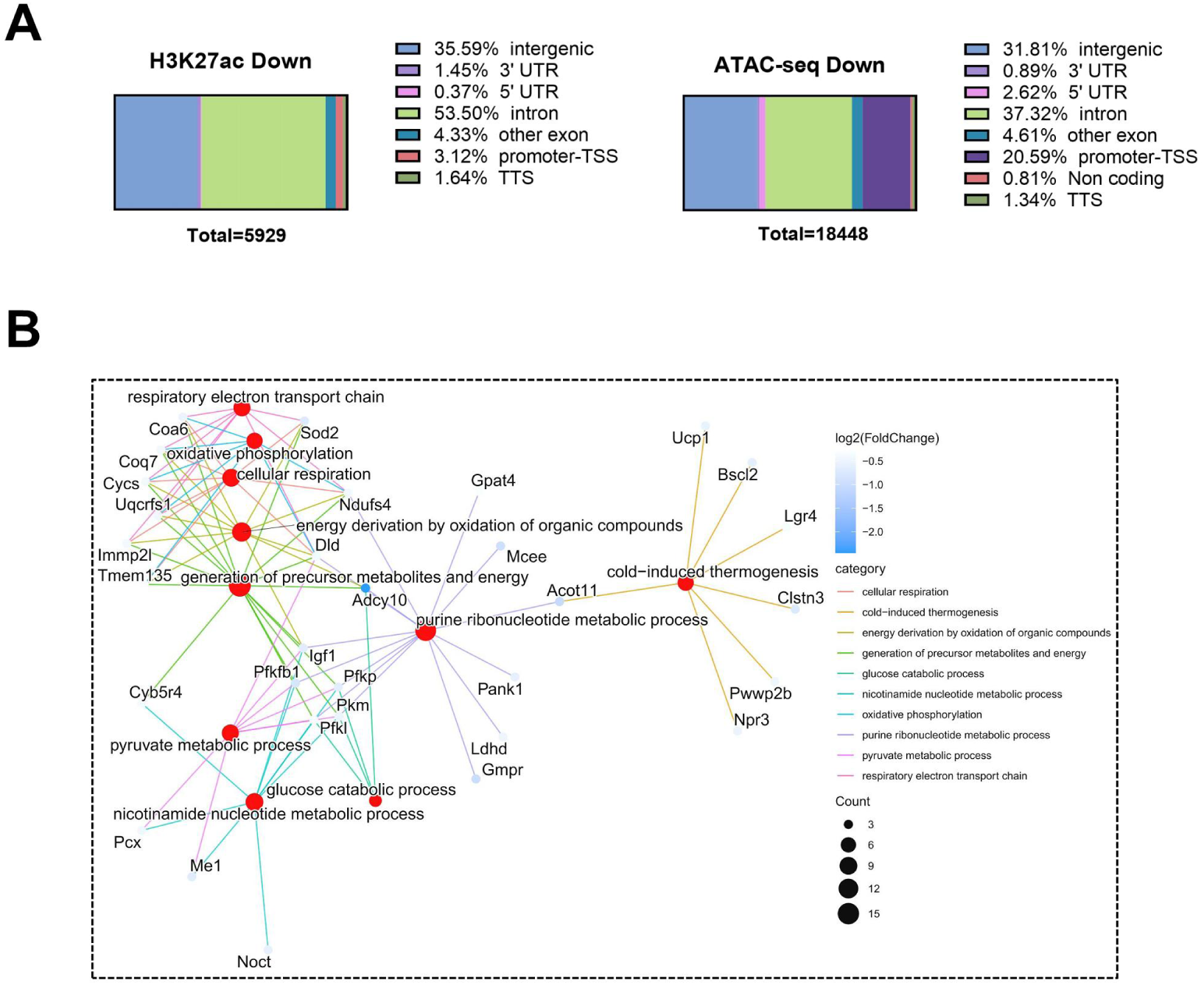
Genomic distribution and functional integration of GPD2-sensitive regulatory elements. (A) Genomic feature annotation of downregulated H3K27ac CUT&Tag and ATAC-seq peak in BAT of WT and GPD2^AKO^ mice. (B) Cnetplot illustrating genes involved in significantly enriched biological process terms derived from the overlapping gene sets of downregulated DEGs, H3K27ac CUT&Tag, and ATAC-seq data. Differential peaks from H3K27ac CUT&Tag and ATAC-seq were annotated to their nearest genes before overlap analysis.

**Fig. S6.**
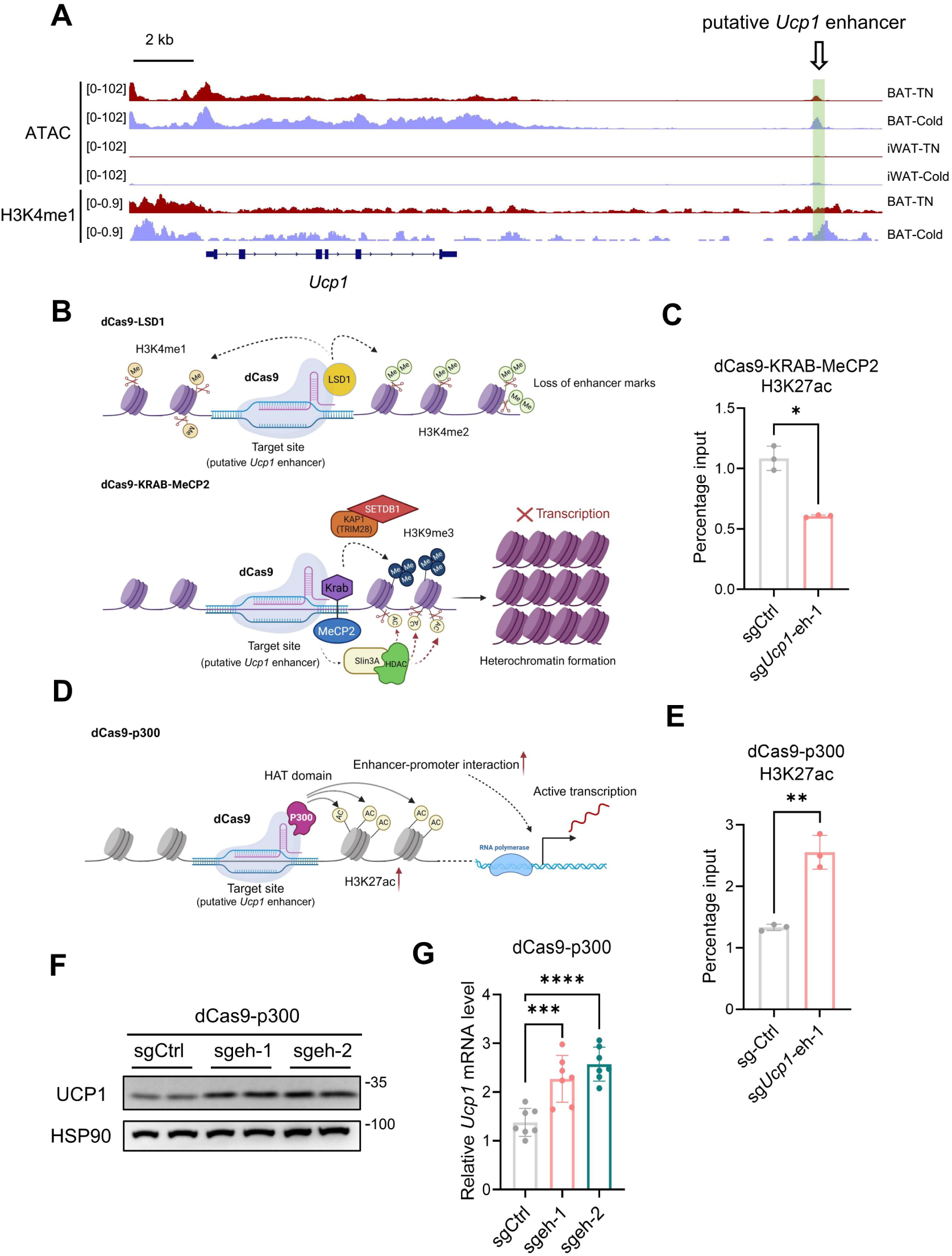
Epigenetic editing of the *Ucp1* enhancer regulates its expression. (A) ATAC-seq (GSE243845) and H3K4me1 (GSE200651) signal intensities at putative *Ucp1* enhancer loci in adipose tissues under TN or cold exposure conditions. (B) Schematic diagrams illustrating the principles of CRISPRi. The dCas9-LSD1 removes enhancer-associated active histone marks including H3K4me1 and H3K4me2. The dCas9-KRAB-MeCP2 system promotes H3K9me3 and reduced H3K27ac enrichment at target site, thereby inducing heterochromatin formation. (C) ChIP-qPCR validation of H3K27ac levels at putative *Ucp1* enhancer loci in the dCas9-KRAB-MeCP2 model. Data were analyzed using two-tailed unpaired Student’s t-test with Welch’s correction. (D) Schematic diagrams illustrating the principles of CRISPRa. The dCas9-p300 system enhance H3K27ac at target site. (E) ChIP-qPCR validate H3K27ac levels at putative *Ucp1* enhancer loci in the dCas9-p300 model. Data were analyzed using two-tailed unpaired Student’s t-test. (F-G) Western blot (F) and qPCR (G) analyses showing UCP1 protein and mRNA levels, respectively, following dCas9-p300-mediated epigenetic editing. Data in (G) were analyzed using ordinary one-way ANOVA with Bonferroni’s multiple comparisons test. *, p < 0.05; **, p < 0.01; ***, p < 0.001; ****, p < 0.0001.

**Fig. S7.**
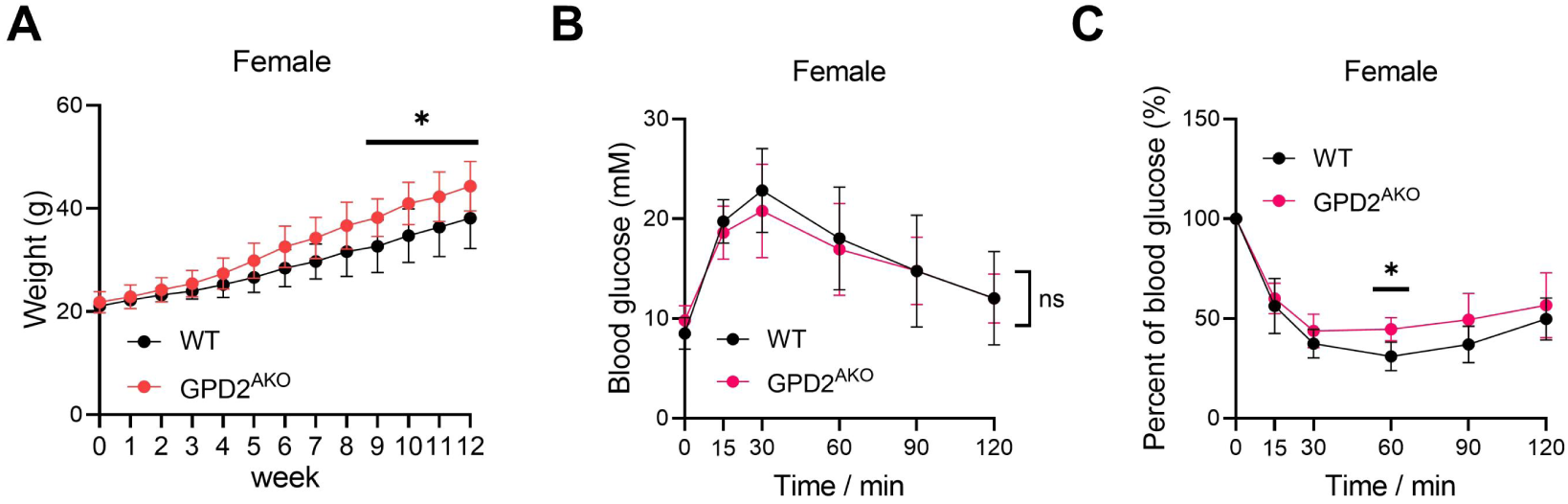
GPD2^AKO^ mice exhibit metabolic dysfunction. (A) Body weight changes in female WT and GPD2^AKO^ mice fed an HFD from 8 weeks of age (n = 7-9). Data were analyzed using two-way ANOVA followed by Bonferroni’s multiple comparisons test. (B-C) GTT (B) and ITT (C) in female WT and GPD2^AKO^ mice fed with HFD (n = 8-9). Data were analyzed using two-way ANOVA followed by Bonferroni’s multiple comparisons test. *, p < 0.05.

**Fig. S8.**
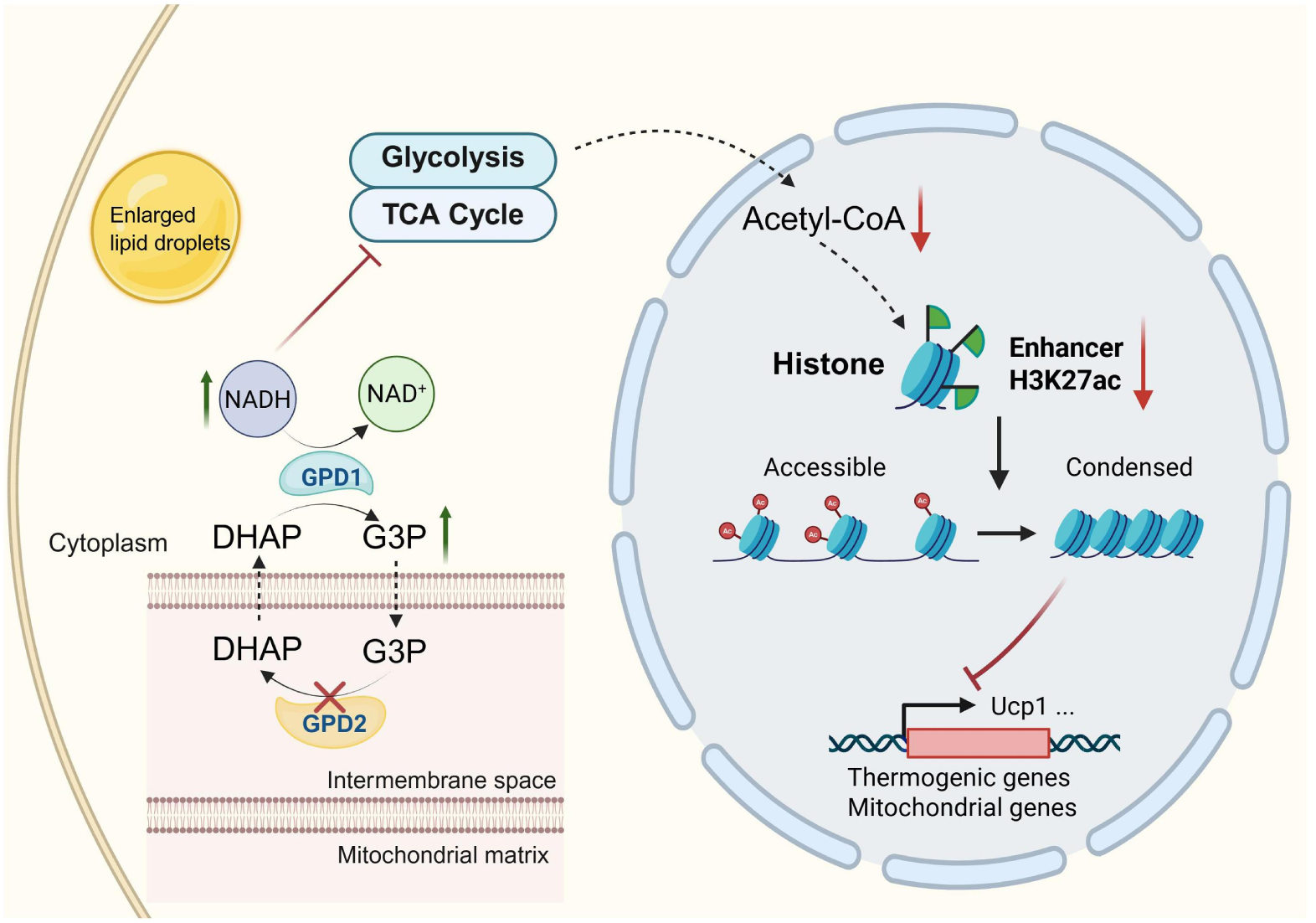
Proposed model illustrating the function of GPD2 in brown adipocytes. Adipose GPD2 expression is positively associated with metabolic homeostasis. Deficiency of GPD2 leads to G3P accumulation. Elevated G3P, the product of GPD1, inhibits GPD1 activity and promotes NADH accumulation. Increased cytosolic NADH suppresses glycolysis and reduces acetyl-CoA levels, thereby diminishing activating histone marks such as H3K27ac at thermogenic gene loci, thereby leading to transcriptional silencing of these genes. Consequently, GPD2^AKO^ mice exhibit impaired energy balance and systemic metabolic dysfunction. The graph model was created with biorender.com.

**Table S1.**
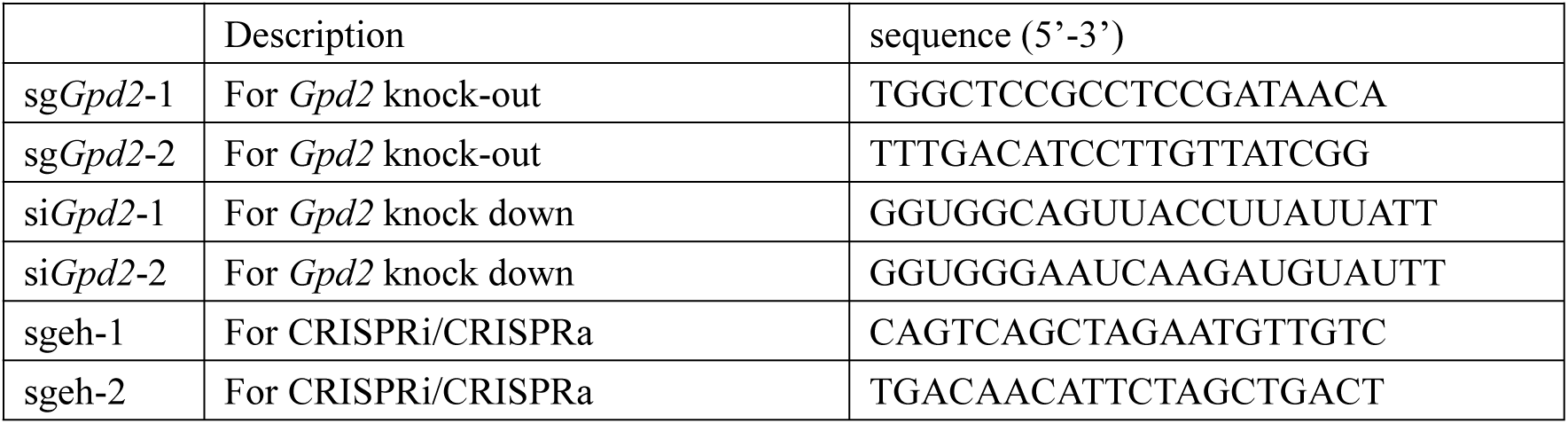
Oligonucleotide sequences used in this study.

**Table S2.**
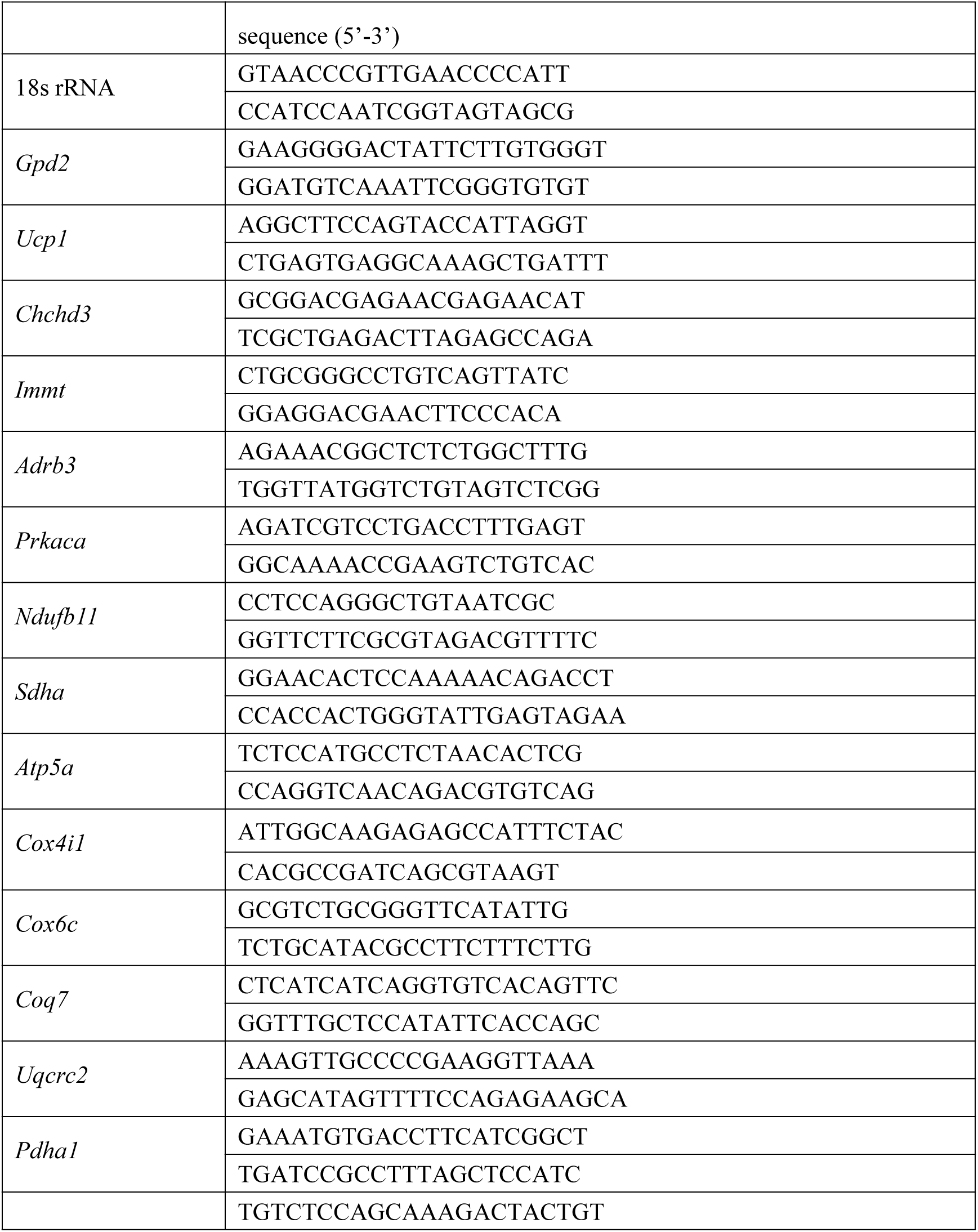
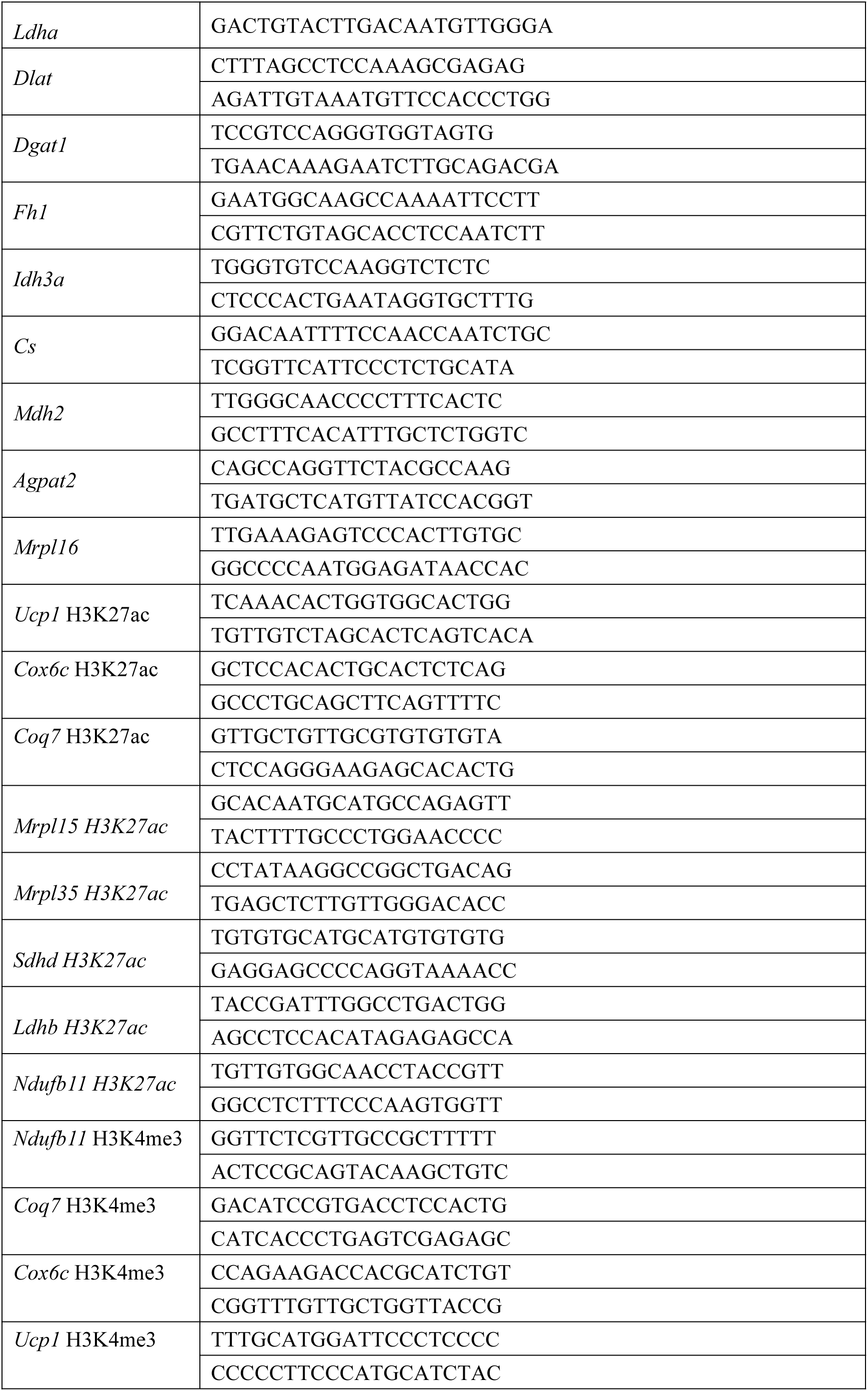
Primer sequence used for qPCR in this study.

**Table S3.** Antibodies used in this study.

|  |  |
| --- | --- |
| GPD2 | Proteintech #17219-1-AP |
| UCP1 | Abcam #ab10983 |
| CHCHD3 | ABclonal #A8584 |
| CHCHD6 | Proteintech #20639-1-AP |
| IMMT | ABclonal #A2751 |
| H3K27ac | ABclonal #A7253 |
| H3K18la | PTMBIO #PTM-7480 |
| H3K27me3 | ABclonal #A22006 |
| H3K4me1 | ABclonal #A22078 |
| Lamin B1 | Proteintech #12987-1-AP |
| FLAG | Proteintech #20543-1-AP |
| ATGL | CST #2439S |
| VDAC-1 | CST #4661S |
| Alpha-Tubulin | Proteintech #11224-1-AP |
| HSP90$\alpha/\beta$ | SANTA #sc-13119 |
| Peroxidase AffiniPure Goat Anti-Rabbit IgG (H+L) | Jackson ImmunoResearch Labs #111-035-003 |
| Peroxidase-AffiniPure Goat Anti-Mouse IgG (H+L) | Jackson ImmunoResearch Labs #115-035-062 |

